# Orchestration of *Drosophila* post-feeding physiology and behavior by the neuropeptide leucokinin

**DOI:** 10.1101/355107

**Authors:** Meet Zandawala, Maria E. Yurgel, Sifang Liao, Michael J. Texada, Kim F. Rewitz, Alex C. Keene, Dick R. Nässel

**Affiliations:** Department of Zoology, Stockholm University, S-10691 Stockholm, Sweden; Department of Biological Sciences, Florida Atlantic University, Jupiter, FL 33458, USA; Department of Biology, University of Copenhagen, Universitetsparken 15, 2100 Copenhagen, Denmark.

**Keywords:** GPCR, insulin signaling, stress resistance, metabolic rate, locomotor activity, neuronal circuit

## Abstract

Behavior and physiology are orchestrated by neuropeptides acting as neuromodulators and/or circulating hormones. A central question is how these neuropeptides function to coordinate complex and competing behaviors. The neuropeptide leucokinin (LK) modulates diverse functions, including circadian rhythms, feeding, water homeostasis, and sleep, but the mechanisms underlying these complex interactions remain poorly understood. Here, we delineate the LK circuitry that governs homeostatic functions that are critical for survival. We found that impaired LK signaling affects diverse but coordinated processes, including regulation of stress, water homeostasis, locomotor activity, and metabolic rate. There are three different sets of LK neurons, which contribute to different aspects of this physiology. We show that the calcium activity of abdominal ganglia LK neurons (ABLKs) increases specifically following water consumption, but not under other conditions, suggesting that these neurons regulate water homeostasis and its associated physiology. To identify targets of LK peptide, we mapped the distribution of the LK receptor (*Lkr*), mined brain single-cell transcriptome dataset for genes coexpressed with *Lkr*, and utilized trans-synaptic labeling to identify synaptic partners of LK neurons. *Lkr* expression in the brain insulin-producing cells (IPCs), gut, renal tubules and sensory cells, and the post-synaptic signal in sensory neurons, correlates well with regulatory roles detected in the *Lk* and *Lkr* mutants. Furthermore, these mutants and flies with targeted knockdown of *Lkr* in IPCs displayed altered expression of insulin-like peptides (DILPs) in IPCs and modulated stress responses. Thus, some effects of LK signaling appear to occur via DILP action. Collectively, our data suggest that the three sets of LK neurons orchestrate the establishment of post-prandial homeostasis by regulating distinct physiological processes and behaviors such as diuresis, metabolism, organismal activity and insulin signaling. These findings provide a platform for investigating neuroendocrine regulation of behavior and brain-to-periphery communication.

## Introduction

Neuropeptides and peptide hormones commonly act on multiple targets in an organism, and for a given neuropeptide these targets can be synchronized and thus orchestrate a specific physiological adaptation or behavior [1–3]. In other cases, the action of a specific neuropeptide can be dissociated in time and space, and therefore occur in a distributed fashion in different circuits of the nervous system [1,2,4]. It can be assumed that peptides expressed in smaller sets of neuroendocrine cells are more likely to serve broad orchestrating functions [4,5]. To explore this assumption we investigated signaling mediated by the neuropeptide leucokinin (LK), which is produced by a small set of neurons and neurosecretory cells in *Drosophila* [6,7].

A central question in biology is how homeostatically regulated behaviors and physiological processes critical for survival interact. LK is an excellent candidate as a factor orchestrating these regimes because it has been implicated in multiple homeostatically regulated functions, including sleep, feeding and response to ionic stress. Previous *in vitro* work has suggested that one of the main functions of LK in adult *Drosophila*, and several other insect species, is to regulate fluid secretion in the Malpighian (renal) tubules (MTs), and, thus, to play an important role in water and ion homeostasis [8–12]. More recently, additional LK functions have been inferred from genetic experiments *in vivo*, such as roles in organismal water retention, survival responses to desiccation and starvation, subtle regulation of food intake, and chemosensory responses [13–18]. Furthermore, it was shown that diminished LK signaling results in an increase in postprandial sleep [19] and impaired locomotor activity [20]. Hence, while LK is critical for behavioral and physiological homeostasis, it is not clear how a relatively small population of neurons can mediate different responses to environmental perturbation. Moreover, it remains unclear whether the different functions revealed are all part of a global orchestrating role of LK in which central and peripheral actions are coordinated at different levels.

To identify broad coordinating actions of LK signaling we generated novel *Lk* and *Lkr* mutant flies. By testing these mutants in various feeding-related physiological and behavioral assays, we found that LK signaling regulates water homeostasis and associated stress, locomotor activity and metabolic rate. From these data, we propose that the regulatory roles of LK can be linked to the orchestration of post-feeding physiology and behavior. One set of LK neurons, the abdominal ganglion LK neurons (ABLKs), but not the ones in the brain, display increased calcium activity in response to rehydration following desiccation. Next, to reveal novel targets of LK peptide, we mapped the distribution of *Lkr* expression. Using two independent *Lkr-GAL4* lines to drive GFP, we show that *Lkr* is expressed in various peripheral tissues, including the gut, Malpighian tubules and sensory cells, which correlates well with the functions suggested by the mutant analysis. In addition, the expression of the *Lkr* in the insulin-producing cells (IPCs) and the phenotypes seen after targeted receptor knockdown indicate interaction between LK and insulin signaling. Thus, the three different types of LK neurons orchestrate post-prandial physiology by acting on different targets in the CNS, as well as renal tubules and intestine.

## Results

### Generation and analysis of *Lk* and *Lkr* mutant flies

To investigate the role of Lk signaling in orchestrating physiology and behavior, we utilized CRISPR/Cas9 to generate GAL4 knock-in mutants for *Lk* and *Lkr* (Fig. 1A). First, we tested the efficiency of the *Lk* and *Lkr* mutants by quantitative real-time PCR (qPCR) and immunolabeling. In qPCR experiments, we found an 80% diminishment of *Lk* expression, whereas *Lkr* mRNA was reduced by about 60% (Fig. 1C). In the homozygous *Lk* mutants, LK immunolabeling is completely abolished in all cells of the CNS (Fig. 1B and D), confirming the efficacy of gene-edited mutants for *Lk and Lkr*. Next, to determine whether there is feedback between components of the LK signaling system we measured LK expression in *Lkr* mutant flies. LK immunolabeling was elevated in abdominal LK neurons (ABLKs) (Fig. 2A and B) and the cell bodies of these neurons were also enlarged (Fig. 2C) probably due to the increased peptide production [see [21]]. Interestingly, the LK immunolabeling in the lateral horn LK (LHLK) neurons of the brain does not change in *Lkr* mutant flies (Fig. 2D and E). Thus, LK levels are differentially regulated in neurons of the brain and the abdominal ganglion, and there appears to be a feedback between receptor and peptide expression in abdominal ABLK neurons of *Lkr* mutant flies.

**Figure 1:**
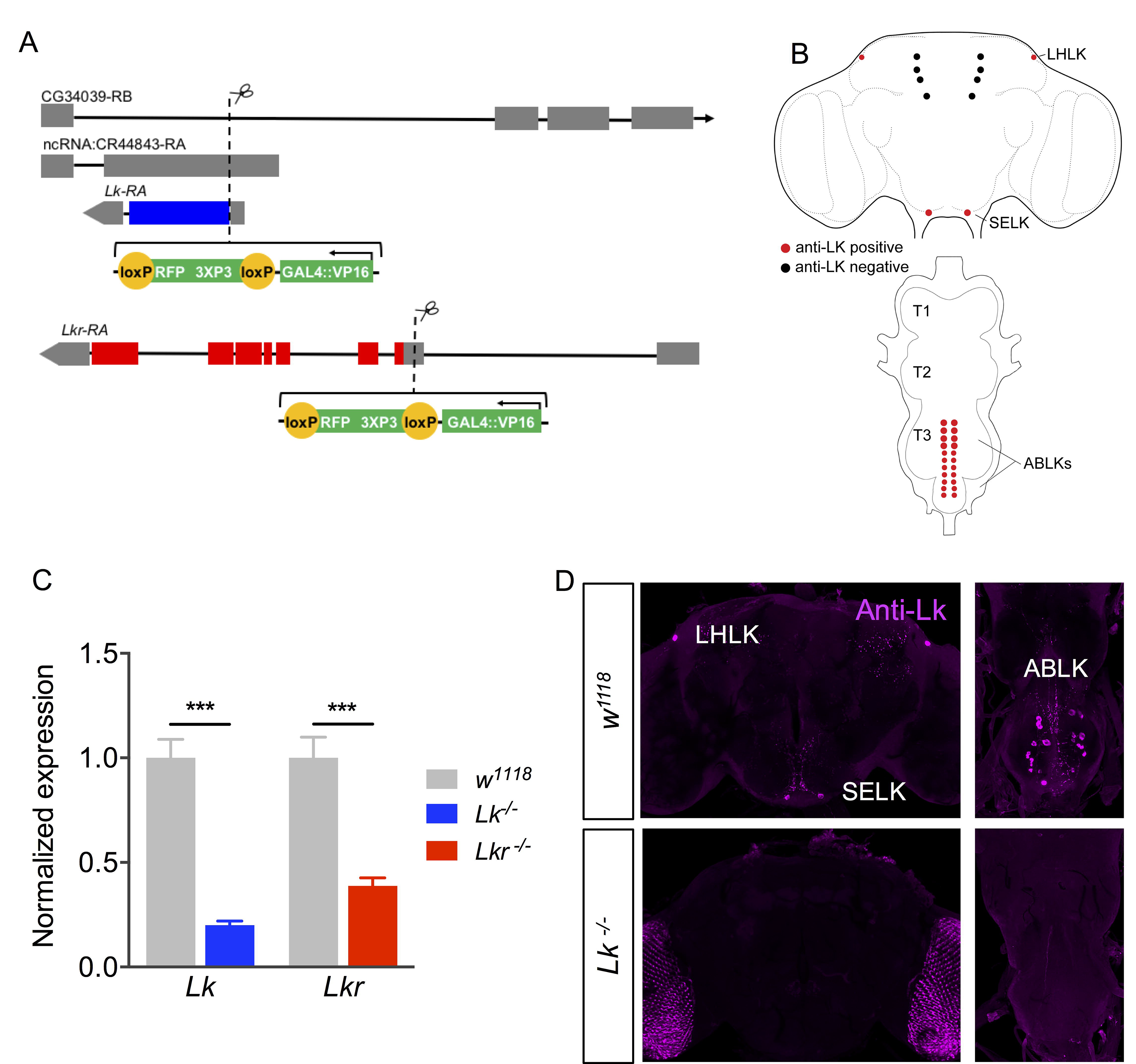
Generation of *Lk* and *Lkr GAL4* knock-in mutants. **(A)** Schematics of the *Lk* and *Lkr* gene loci and the locations of construct insertion to generate *GAL4* knock-in mutants. **(B)** A schematic of the adult CNS showing the location of LK-expressing neurons [based on [6,7,17]]. LHLK, lateral horn LK neuron; SELK, subesophageal ganglion LK neuron; ABLK, abdominal LK neuron, T1 - T3, thoracic neuromeres. **(C)** Quantitative PCR shows a significant reduction in *Lk* and *Lkr* transcripts in *Lk* and *Lkr* homozygous mutants, respectively. (*** p < 0.001 as assessed by unpaired *t* test). **(D)** LK-immunoreactivity is completely abolished in the brain and ventral nerve cord of *Lk* mutants.

**Figure 2:**
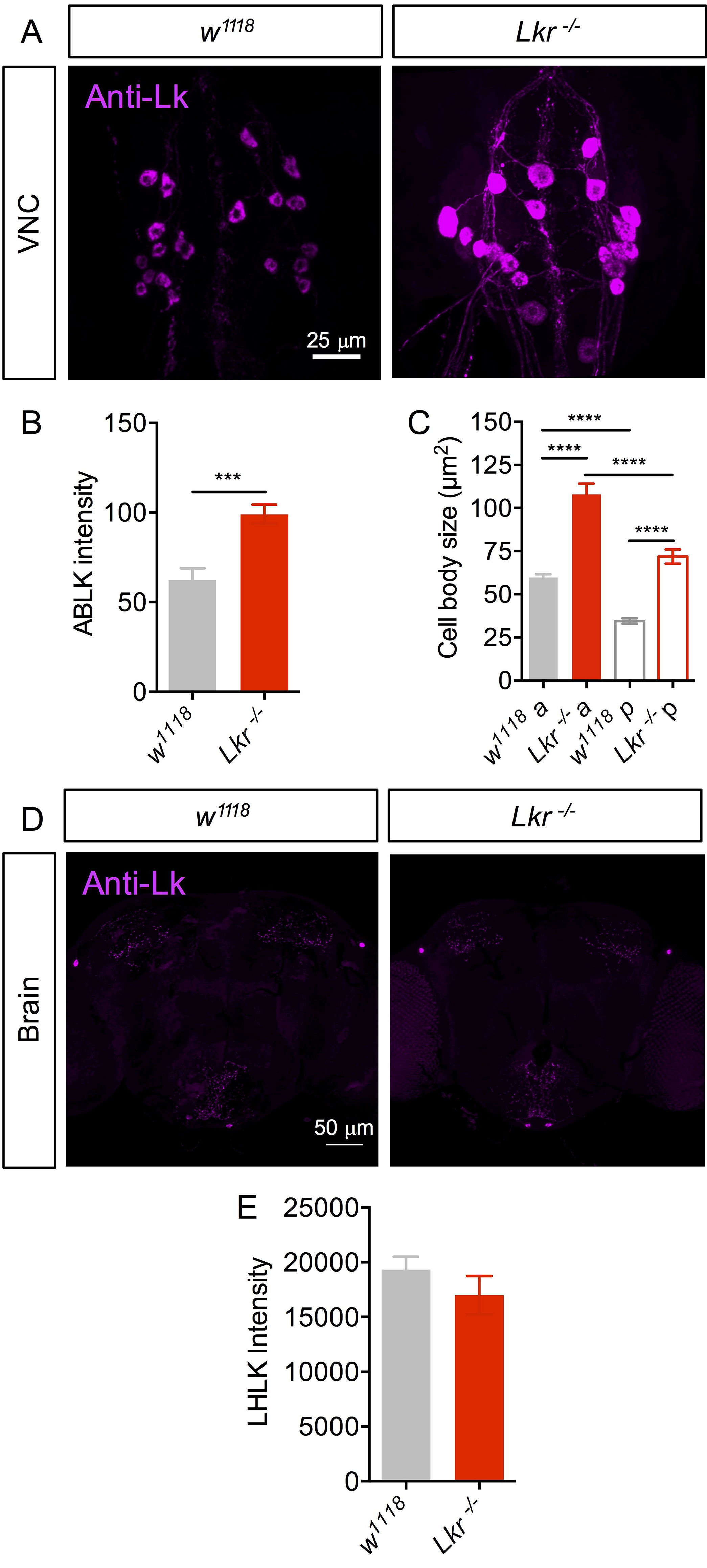
LK cell body size and peptide levels in *Lkr* mutants. **(A)** LK-immunoreactivity in abdominal LK neurons (ABLKs) of *Lkr* mutant and control flies. **(B)** Staining intensity and **(C)** cell body size of both the anterior (a) and posterior (p) ABLKs is increased in *Lkr* mutants compared to control flies. **(D)** LK-immunoreactivity in brain lateral horn LK neurons (LHLKs) of *Lkr* mutant and control flies. **(E)** The intensity of LK staining is unaltered in *Lkr* mutants. (**** p < 0.0001 as assessed by one-way ANOVA followed by Tukey’s multiple comparisons test for **C** and *** p < 0.001 as assessed by unpaired *t* test for **B**).

Previous studies have demonstrated the role of LK signaling in MT secretion [9,12] and a possible secondary effect of this on desiccation and starvation resistance [14,16,17]. We therefore tested survival of *Lk* and *Lkr* mutant flies maintained under desiccation and starvation conditions. Both homozygous and heterozygous *Lk* (*Lk*-GAL4^CC9^) and *Lkr* mutants (*Lkr*-GAL4^CC9^), survived longer under conditions of desiccation and starvation (Fig. 3A-D). To determine whether changes in water content contributed to these survival differences, we assayed flies for their water content under normal conditions and after 9 hours of desiccation. As expected, *Lk* and *Lkr* mutant flies displayed higher water content than control flies under normal conditions as well as after desiccation (Fig. 3E). Therefore, loss of Lk or Lkr promotes water retention and improves survival under desiccation conditions.

**Figure 3:**
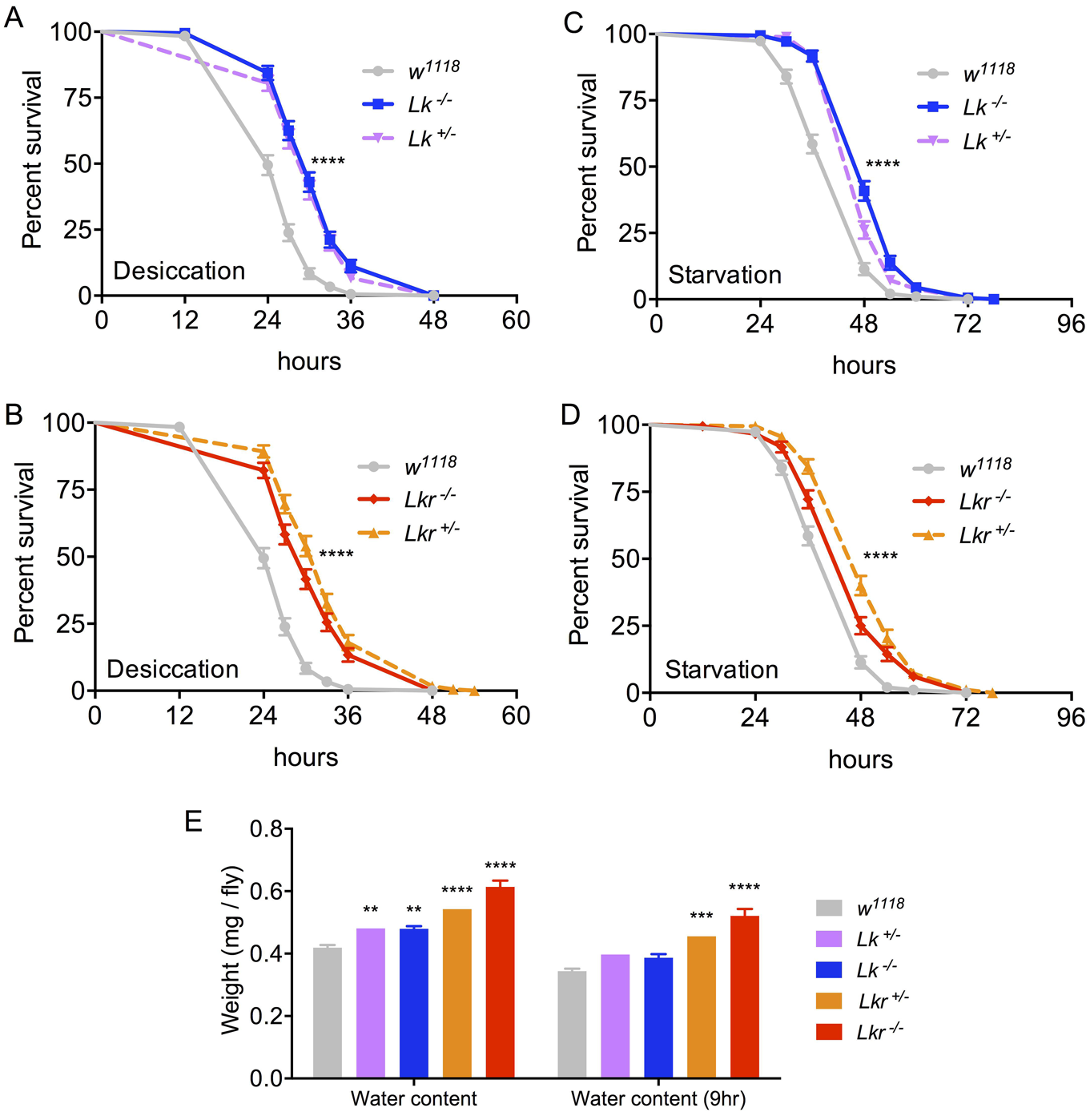
*Lk* and *Lkr* mutants have altered stress resistance and water content. Survival under desiccation is increased in both **(A)** *Lk* and **(B)** *Lkr* mutants. Survival under starvation is also increased in both **(C)** *Lk* and **(D)** *Lkr* mutants. Data are presented in survival curves and the error bars represent standard error (**** p < 0.0001, as assessed by Log-rank (Mantel-Cox) test). **(E)** Hydrated and 9-hour-desiccated (9 h) *Lk* and *Lkr* mutant flies show increased water content compared to control flies. (** p < 0.01, *** p < 0.001, **** p < 0.0001 as assessed by one-way ANOVA followed by Tukey’s multiple comparisons test).

To determine which of the LK neurons respond to starvation, desiccation and/or water ingestion we monitored the calcium activity of LK neurons using the CaLexA system [22]. We found that only the ABLKs, but not the LK neurons in the brain (not shown), were activated following re-watering (drinking) (Fig. 4A). The activation of ABLKs can be seen as increased GFP intensity as well as the number of cells that could be detected (Fig. 4B and C). Moreover, these cells did not display activation when the flies are placed under starvation, desiccation or on artificial diet. These results further support the role of ABLKs in the regulation of water homeostasis.

**Figure 4:**
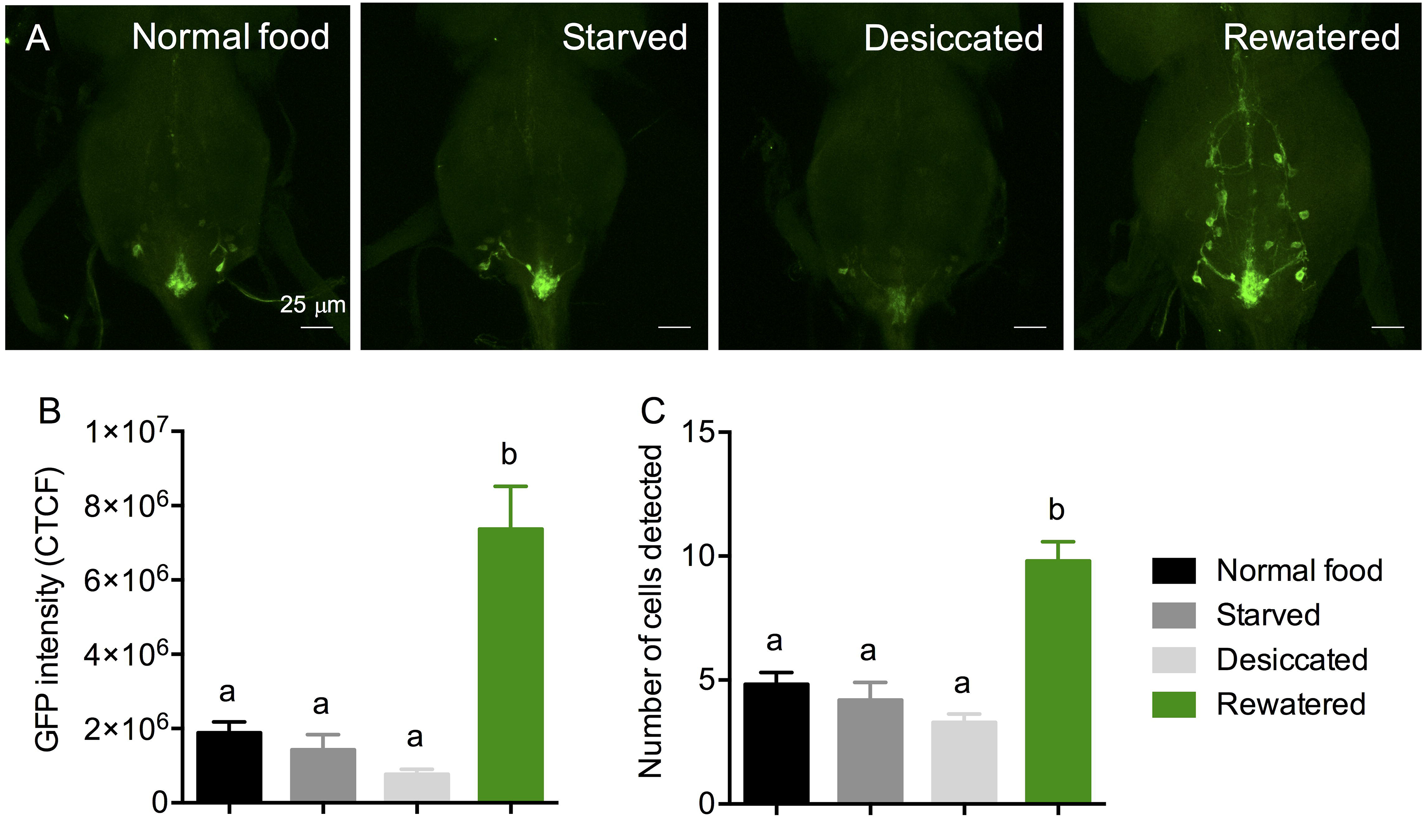
Calcium activity of ABLKs under nutritional and osmotic stress. **(A)** The calcium activity of ABLKs, as measured using CaLexA [22], is low in flies that have been starved, desiccated, or incubated on normal artificial food but increased in flies that have been rewatered (desiccated and then incubated on 1% agar). **(B)** The GFP intensity of ABLKs is increased in rewatered flies compared to other conditions. **(C)** The number of ABLKs that could be detected is higher in rewatered flies compared to other conditions. (assessed by one-way ANOVA followed by Tukey’s multiple comparisons test).

Next, we tested the *Lk* and *Lkr* mutants for the strength of the proboscis extension reflex (PER) under different sucrose concentrations (Fig. 5A-D and Supplementary Table 1). The *Lk* mutant flies displayed a reduced PER (Fig. 5C) and this phenotype was rescued by UAS-*Lk* in the homozygous *GAL4*-insertion mutants (Fig. 5D). This reduction in PER was also seen after inhibition of LK neurons by targeted expression of UAS-*TNT* (Fig. 5B). However, the *Lkr* mutant flies displayed the opposite behavior, showing increased PER that could also be rescued by UAS-*Lkr* expression (Fig. 5A). Finally, we used an assay for short term feeding (over 30 min), in which the amount of ingested blue-dyed food was measured in fly homogenates. In this assay, there was no difference in food intake between mutant flies and controls, either in starved or fed conditions (Fig. 5E). This lack of effect was also seen when the LK neurons were inhibited by targeted expression of UAS-*TNT* (Fig. 5F). Therefore, LK neurons seem to regulate the propensity of animals to initiate reflexive feeding, without affecting total meal volume.

**Figure 5:**
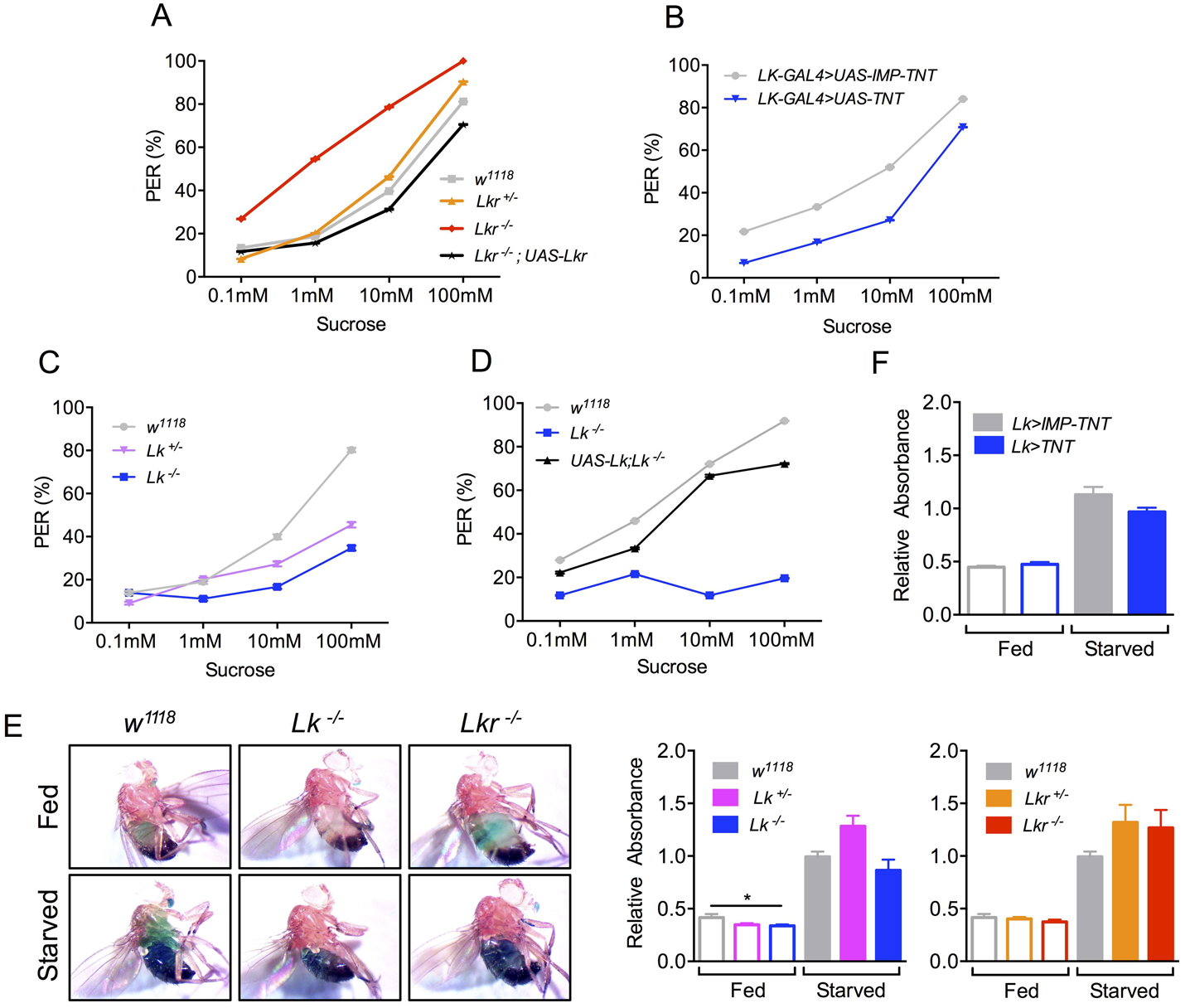
*Lk* and *Lkr* mutants show varying phenotypes in different feeding assays. **(A)** *Lkr* mutants show increased motivation to feed in proboscis extension reflex (PER) which could be rescued to control levels by driving *UAS-Lkr* with *Lkr-GAL4*^*CC*9^. **(B)** Interestingly, targeted expression of tetanus toxin (to block synaptic transmission) in *Lk* neurons using *Lk-GAL4* caused a decrease in PER. **(C)** Both the homozygous and heterozygous *Lk* mutants also show decreased PER and this phenotype could be rescued in **(D)** the homozygous flies. See Supplementary Table 1 for the statistics of graphs A-E. **(E)** Starved and fed *Lk* and *Lkr* mutants do not show any differences in short-term feeding compared to control flies as measured using a blue-dye feeding assay (assessed by one-way ANOVA). **(F)** Expression of tetanus toxin in *Lk* neurons also has no effect on short-term feeding.

Activity and metabolic rate are acutely regulated by food availability and environmental stress. To determine whether LK regulates these processes we simultaneously recorded animal activity and metabolic rate using stop-flow indirect calorimetry [23]. Single *Lk* and *Lkr* mutant flies were tested for locomotor activity and metabolic rate (vCO_2_) over a 24-hour period. The *Lk* mutants displayed reduced locomotor activity, with homozygotes displaying almost no morning or evening activity peaks (Fig. 6A and B). The metabolic rate of these mutant flies was also reduced over the entire period of observation (Fig. 6C and D). The *Lkr* mutants displayed a similar reduction in both locomotor activity and metabolic rate, except that the heterozygotes displayed no change in locomotor activity (Fig. 6E-H). We also used the standard *Drosophila* activity monitor system (DAMS) to verify our locomotor activity results from the above setup. Indeed, we obtained similar results to those above, with *Lk* and *Lkr* mutants displaying reduced activity (Fig. S1A and B). Together, these findings suggest that LK stimulates both metabolic rate and activity.

**Figure 6:**
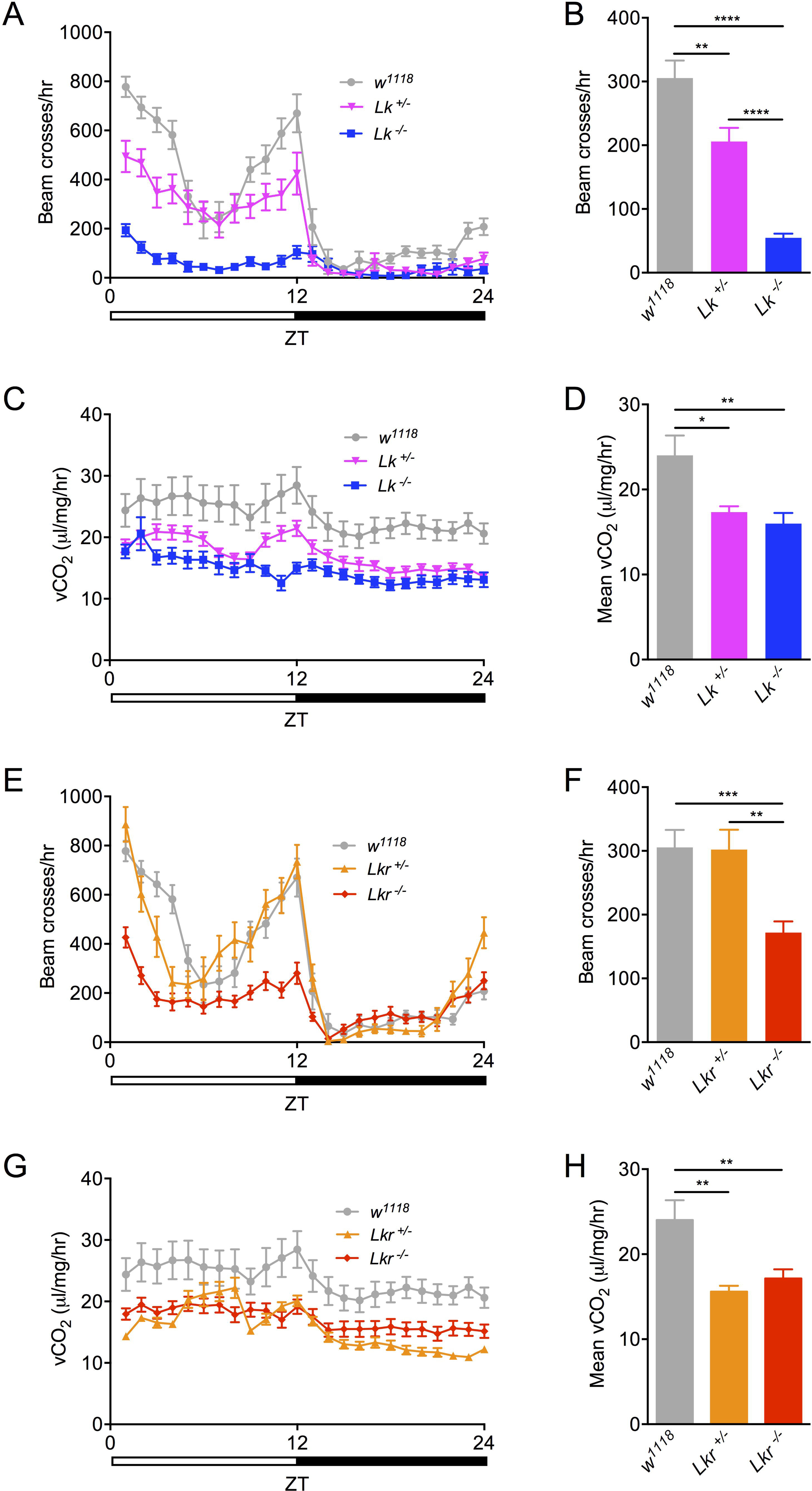
Total activity and metabolic rate is lowered in individual *Lk* and *Lkr* mutants. **(A)** Locomotor activity pattern of individual *Lk* homozygous and heterozygous mutants measured over 24 hours. **(B)** Total locomotor activity of *Lk* mutants is lowered compared to control flies. **(C)** Metabolic rate rhythms of individual *Lk* homozygous and heterozygous mutants measured over 24 hours. **(D)** Average metabolic rate of *Lk* mutants is lowered compared to control flies. **(E)** Locomotor activity pattern of individual *Lkr* homozygous and heterozygous mutants measured over 24 hours. **(F)** Total locomotor activity of *Lkr* mutants is lowered compared to control flies. **(G)** Metabolic rate rhythms of individual *Lkr* homozygous and heterozygous mutants measured over 24 hours. **(H)** Average metabolic rate of *Lkr* mutants is lowered compared to control flies. (* p < 0.05, ** p < 0.01, *** p < 0.001, **** p < 0.0001 as assessed by one-way ANOVA).

### Identifying targets of LK

The expression of *Lk* and *Lkr* in the central nervous system (CNS) and periphery raises the possibility that distinct populations or neural circuits regulate different behaviors. The *Lk* and *Lkr GAL4* knock-in mutants (*GAL4*^*CC*9^) that we generated using CRISPR/Cas9 enable simultaneous knockdown and visualization of the distribution of peptide and receptor gene expression in different tissues. Since the *GAL4* is inserted within the gene itself, the retention of all the endogenous regulatory elements should in theory allow *GAL4* expression to mimic that of the *Lk* and *Lkr*. Thus, the *Lk-GAL4*^*CC*9^ expression observed (Fig. S2) is very similar to that seen in earlier reports using conventional *Lk-GAL4* lines [7,18]. With a few exceptions, the pattern of *Lk-GAL4*^CC9^ expression also matches that of LK immunolabeling (Fig. S2C and D). Notably, a set of 5 pairs of GFP-labeled lateral neurosecretory cells does not display LK immunolabeling in third instar larvae or adult flies (Fig. S2C and S3A). These neurons are known as ipc-1 and ipc-2a, and they express ion transport peptide (ITP), short neuropeptide F (sNPF) and *Drosophila* tachykinin (DTK) [24,25].

Since the cellular expression of *Lkr* in *Drosophila* is poorly known we utilized our *Lkr-GAL4*^*CC*9^ line to drive GFP and analyzed CNS and peripheral tissues. We compared the expression of our *Lkr-GAL4*^*CC*9^ to that of another *Lkr-GAL4* (*Lkr-GAL4::p65*) generated using a BAC clone as described previously [26] and found a high degree of overlapping expression patterns between the two drivers. In the periphery, the stellate cells of the MTs express *Lkr-GAL4*^*CC*9^ (Fig. 7A) as expected from earlier work demonstrating functional expression of the Lkr in these cells [9,12]. Furthermore, *Lkr-GAL4*^*CC*9^ driven GFP was detected in endocrine cells of the posterior midgut (Fig. 7B), in the anterior midgut (Fig. 7C and D), and in muscle fibers of the anterior hindgut and rectal pad (Fig. 7E and F). *Lkr-GAL4*^*CC*9^>*GFP* expression was also present in peripheral neurons (Fig. S4A), the dorsal vessel as well as the nerve fibers innervating it (Fig. S4A), and the sensory cells of the legs, mouthparts and anterior wing margin (Fig. S4B-D). In third instar larvae, we could also detect *Lkr-GAL4*^*CC*9^ expression in the stellate cells of the MTs (Fig. S5A and D), in the ureter (Fig. S5A), in muscle fibers of the gastric caeca, midgut and hindgut (Fig. S5A-C), as well as in the endocrine cells of the midgut (Fig. S5B and C). The BAC-engineered *Lkr-GAL4* had a much sparser expression pattern, with GFP detected in stellate cells of larval (Fig. S6A) and adult (Fig. S6D-E) MTs, and in the larval hindgut (Fig. S6B). Interestingly, the shape of the stellate cells in adults varied between cuboidal and the more typical star-shaped morphology (Fig S6C and D).

**Figure 7:**
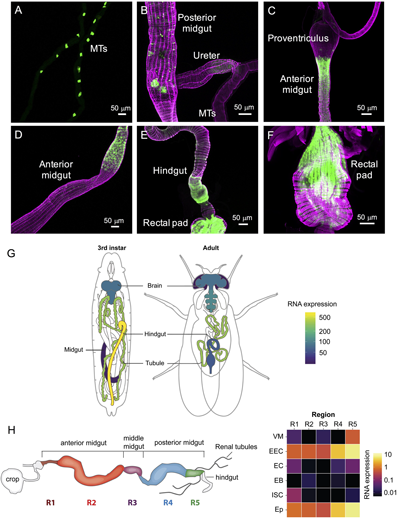
*Lkr* is expressed in the adult gut and Malpighian tubules. *Lkr-GAL4*^*CC*9^ drives GFP (*pJFRC81-10xUAS-Syn21-myr::GFP-p10*) expression in the adult **(A)** stellate cells in Malpighian tubules, **(B)** enteroendocrine cells in the posterior midgut, **(C and D)** anterior midgut, **(E)** hindgut and **(F)** rectal pad. Muscles (F-actin filaments) in all the preparations (except B) have been stained with rhodamine-phalloidin (magenta). Note the expression of GFP in hindgut and rectal pad muscles. **(G)** Schematics of third instar larvae and adult fly showing the expression of *Lkr*. (data from FlyAtlas.org, [27]). **(H)** A schematic of the adult gut and heat map showing expression of *Lkr* in different regions of the gut (R1 to R5) and its various cell types (VM, visceral muscle; EEC, enteroendocrine cell; EC, enterocyte; EB, enteroblast; ISC, intestinal stem cell; Ep, epithelium. Data was mined using Flygut-*seq* [28].

In general, the expression of the new *Lkr-GAL4*^*CC*9^ line is in agreement with the BAC/promoter fusion line and available immunolabeling data, suggesting that they largely recapitulate the endogenous receptor expression pattern. To further validate the authenticity of the GFP expression in the periphery, we examined *Lkr* expression in two publicly available resources for gene expression, FlyAtlas [27] and Flygut-seg [28]. FlyAtlas reveals that *Lkr* is expressed in the larval and adult hindgut, MTs and CNS (Fig. 7G). Moreover, the Flygut-*seg* data base shows that *Lkr* is expressed in enteroendocrine cells of the midgut, in visceral muscles near the hindgut and in the gut epithelium [28] (Fig. 7H). Thus, the transcript expression data correlate well with the GAL4 expression pattern.

The expression pattern of *Lkr-GAL4*^*CC*9^ and the *Lkr-GAL4* also matched well within the brain. Both GAL4 lines drive GFP expression in a relatively large number of neurons in the larval (Fig S3B and S7A) and adult CNS (Fig. S7B-C and S8), but we focus here on two sets of identified peptidergic neurons in the brain (Fig. 8). Both, the *Lkr-GAL4*^*CC*9^ and *Lkr-GAL4*, drove GFP expression in the brain IPCs, as identified by anti-DILP2 staining, and in the 5 pairs of brain ipc-1/ipc-2a cells, that colocalize anti-ITP staining (Fig. 8). In addition, comparison to the single-cell transcriptome dataset of the entire *Drosophila* brain [29] identified coexpression between *Lkr* and *DILP2*, *3* and *5*, as well as *Lkr* and *ITP* (Fig. 9). *Lkr* is widely expressed in the *Drosophila* brain with transcripts expressed in cells of various clusters, including the peptidergic cell cluster (marked with *dimm*) and the glia cell cluster (marked with *repo*) (Fig. 9A). Within the peptidergic cell cluster, *Lkr* is coexpressed with *ITP* (Fig. 9B) and in IPCs along with *DILP2, 3* and *5* (Fig. 9C and D). Our receptor expression data further emphasizes the important interplay between LK signaling within the CNS and systemic LK action that targets several peripheral tissues, which together orchestrate physiology and behavior.

**Figure 8:**
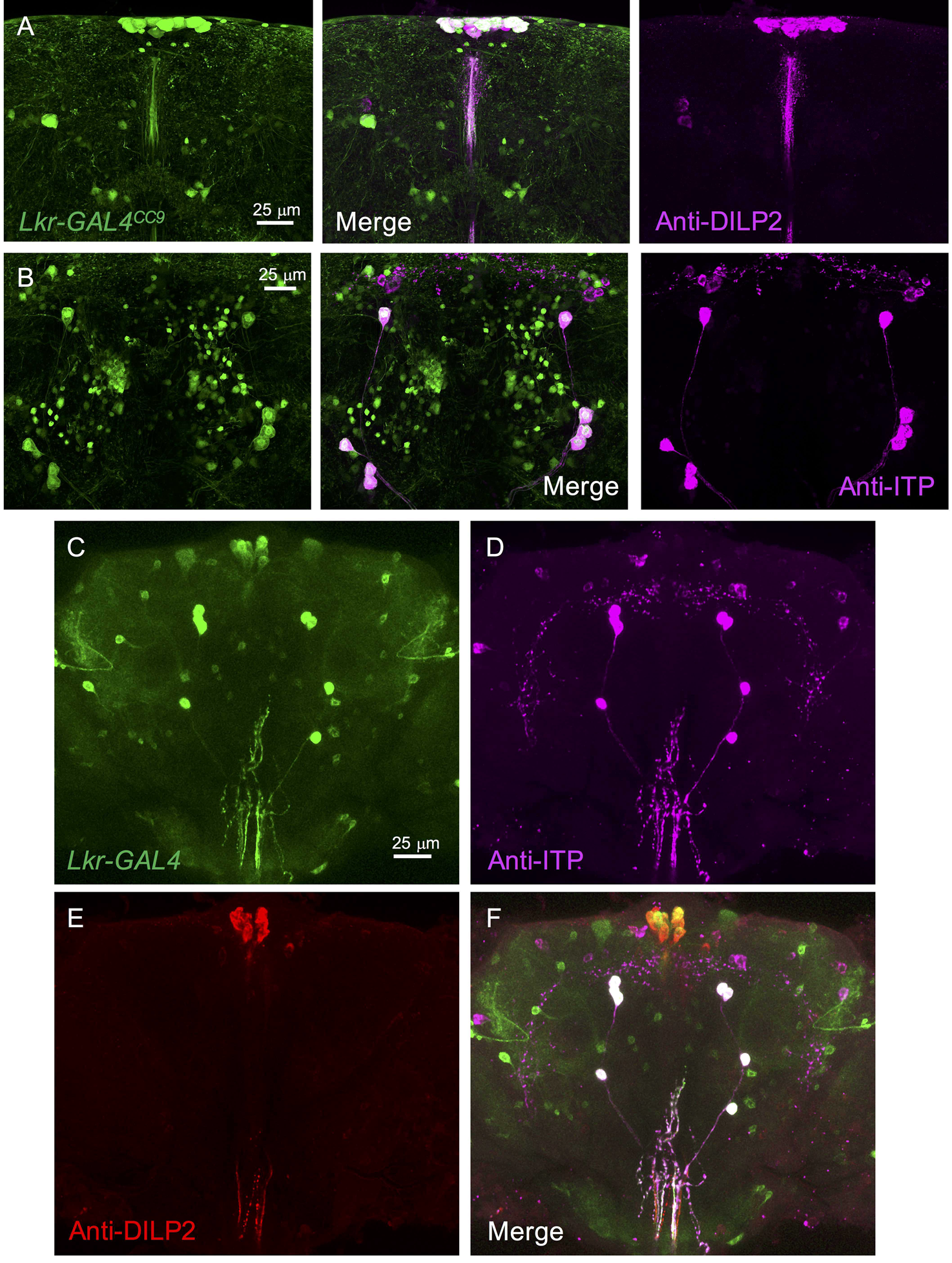
*Lkr* is expressed in identified peptidergic neurosecretory cells of the adult brain. *Lkr-GAL4*^*CC*9^ drives GFP (*pJFRC81-10xUAS-Syn21-myr:: GFP-p10*) expression in **(A)** insulin-producing cells (labeled with anti-DILP2 antiserum) and **(B)** ion transport peptide (ITP)-producing lateral neurosecretory cells in the brain (labeled with anti-ITP antiserum). **(C)** *Lkr-GAL4* drives GFP (*UAS-mCD8GFP*) expression in the adult **(D and F)** ITP-producing cells and **(E and F)** insulin-producing cells.

**Figure 9:**
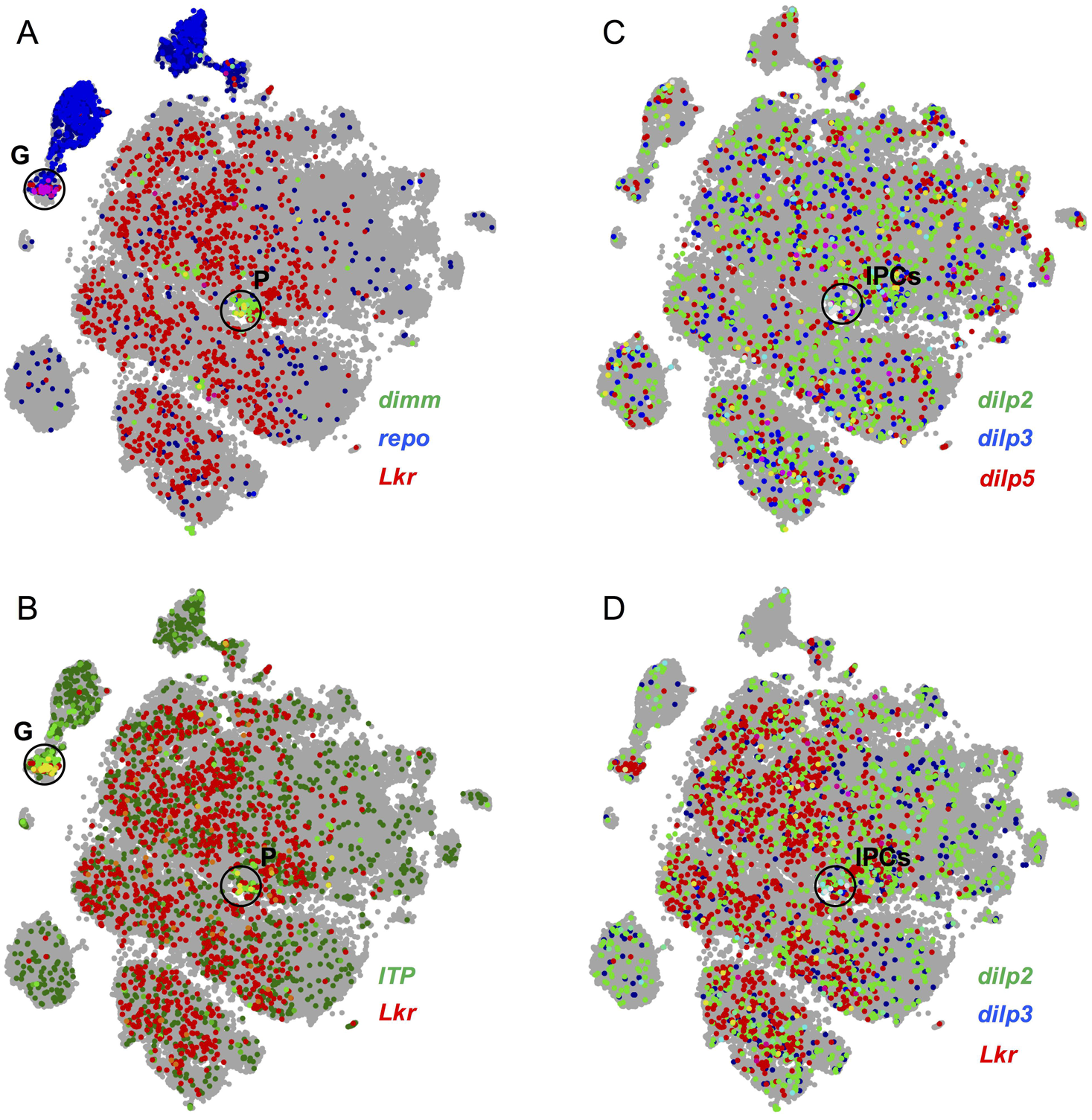
*Lkr* is coexpressed with peptidergic and glial markers. Mining the single-cell transcriptome atlas of the *Drosophila* brain reveals that *Lkr* is coexpressed with **(A)** *repo* (glial marker; cell cluster marked G) and *dimm* (peptidergic cell marker; cell cluster marked P). **(B)** Within both the glial and peptidergic cell clusters, *Lkr* is coexpressed with ITP. Within the peptidergic cell cluster, **(C)** insulin-producing cells expressing *DILP2, 3* and *5* could be identified (cluster marked IPCs), a subset of which express *Lkr* **(D)**. Data was mined using SCope (http://scope.aertslab.org) [29]. In both **(C)** and **(D)**, cells expressing all three genes are colored in white.

To establish the nature of connections (synaptic versus paracrine) between LK neurons and the IPCs, and to identify other neurons downstream of LK signaling, we employed the *trans*-Tango technique for anterograde trans-synaptic labeling of neurons [30]. Using *Lk-GAL4* to drive expression of the system, we see strong GFP-labeling (pre-synaptic marker) in SELK neurons and expression of the post-synaptic marker (visualized by mtdTomato tagged with HA) is seen in several SEG neurons some of which have axons that project to the pars intercerebralis (Fig. 10 A and B). *Lkr* is expressed in the IPCs, which have dendrites in the tritocerebrum and subesophageal zone where the LK post-synaptic signal is found (Fig. S10), so we asked if the IPCs are post-synaptic to SELKs. However, no colocalization is seen between the IPCs and post-synaptic signal of LKs. In addition, the post-synaptic signal is not coexpressed with Hugin neurons (labeled with anti-CAPA antibody) although these have similar axonal projections (Fig. S9). Hence, these anatomical data indicate that the IPCs express the Lk receptor, but may receive non-synaptic (paracrine) inputs from LK neurons, or possibly via the circulation from ABLKs.

**Figure 10:**
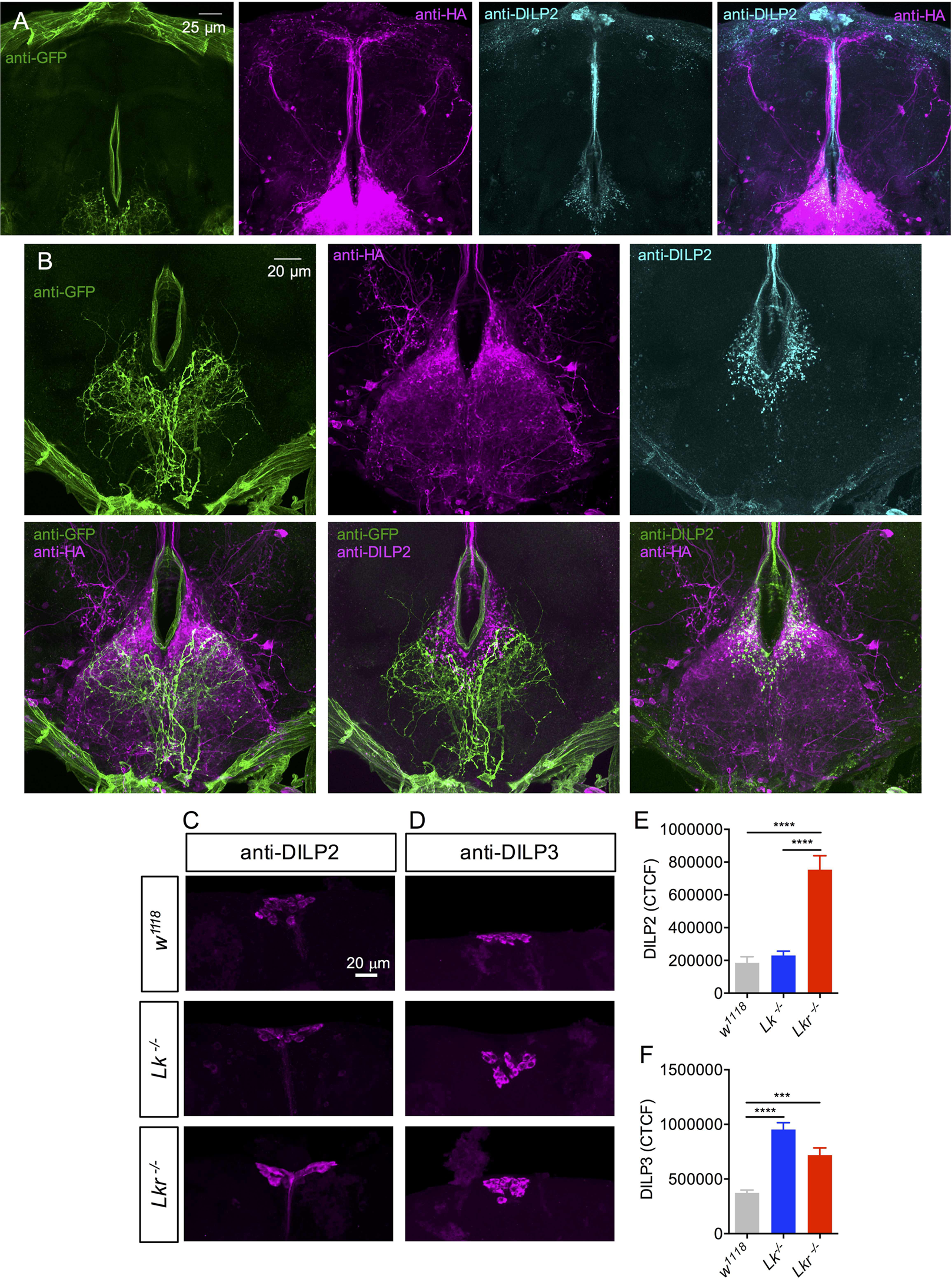
Anatomical and functional Interactions between LK and insulin signaling. **(A)** Expression of *trans*-Tango components [30] using *Lk-GAL4* generates a pre-synaptic signal (labeled with anti-GFP antibody) in the subesophageal ganglion (SEG) and a post-synaptic signal (labeled with anti-HA antibody) in the SEG and pars intercerebralis which does not colocalize with insulin-producing cells or their axons (labeled with anti-DILP2 antibody). **(B)** Higher magnification of the SEG showing the pre-synaptic and post-synaptic signals and the lack of colocalization with anti-DILP2 staining. **(C, E)** *Lkr* homozygous mutants show increased DILP2 immunoreactivity in insulin-producing cells (IPCs) of the adult brain. **(D, F)** Both *Lk* and *Lkr* homozygous mutants show increased DILP3 immunoreactivity in IPCs of the adult brain. (*** p < 0.001, **** p < 0.0001, as assessed by one-way ANOVA followed by Tukey’s multiple comparisons test). CTCF, corrected total cell fluorescence.

Since *Lkr* is expressed in the IPCs we wanted to examine if the expression of DILPs is altered in *Lk* and *Lkr* mutants. In *Lk* mutant flies, DILP3 immunolabeling is increased and in *Lkr* mutants both DILP2 and DILP3 levels are significantly higher (Fig. 10C-F), indicating that LK could affect the release of DILP2 and DILP3 (as increased immunolabeling has been proposed to reflect decreased peptide release [31]). No effect on DILP5 levels was seen for any of the mutants, suggesting that LK selectively modulates DILP function (Fig. S11).

Next, we examined *DILP2*, *DILP3* and *DILP5* transcript levels by qPCR after targeted knockdown of the *Lkr* in the IPCs of flies using two different *Lkr*-RNAi lines and a *DILP2-GAL4* driver; also different diets were tested since *DILP* expression in IPCs are influenced by carbohydrate and protein levels in the food [32]. The experimental flies developed to pupation on normal diet and were transferred as adults to three different diets, high sugar+high protein, low sugar+high protein and normal diet. *UAS-Lkr-RNAi-#1* did not drive efficient knockdown and was thus excluded from the analysis; data shown for *UAS-Lkr-RNAi-#2*. Significant effects on *DILP* transcripts were only seen for *DILP3*, which was increased in flies after *Lkr*-RNAi under normal and high-sugar+high-protein diets, and *DILP5*, which was decreased in normal diet. Having noted an effect on DILP/*DILP* levels in mutant flies and after *Lkr* knockdown in the IPCs we went on to determine the effects of this manipulation on fly weight as well as survival during starvation and desiccation. As seen in Fig. S12, there was a slight increase in survival during desiccation and a small increase in dry weight of the flies with reduced *Lkr* in IPCs.

Taken together, we identify roles for the Lkr within the CNS and in the periphery that uniquely regulate physiological homeostasis. The *Lkr* expression in the periphery suggests LK signaling to be associated with water balance, gut function and chemosensation (Fig. 12). Within the CNS, LK signaling modulates specific neurosecretory cells of the brain that are known to regulate stress responses, feeding, metabolism, energy storage and activity patterns, including sleep (Fig. 12) [24,33–37].

**Figure 11:**
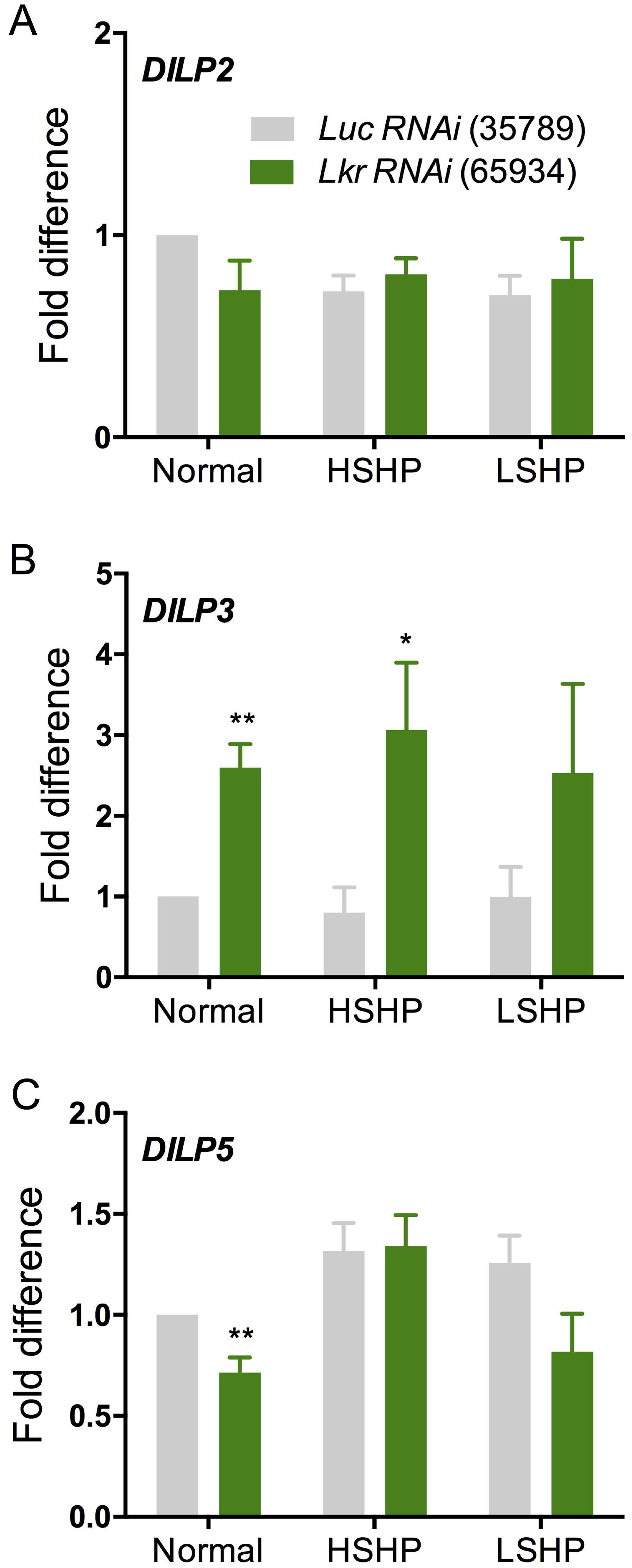
*Lkr* knockdown in insulin-producing cells affects insulin expression. **(A)** Quantitative PCR shows no difference in *DILP2* transcript levels between control flies (*DILP2 > Luciferase-RNAi*) and flies with *Lkr* knockdown in insulin-producing cells (IPCs) that were reared as adults on normal diet, high sugar and high protein diet (HSHP) or low sugar and high protein diet (LSHP). **(B)** *DILP3* transcript levels are upregulated in *DILP2* > *Lkr-RNAi-#2 (BL65934)* flies reared on normal and HSHP diets. **(C)** *DILP5* transcript is downregulated in *DILP2 > Lkr-RNAi-#2 (BL65934)* flies reared on normal diet. (* p < 0.05 and ** p < 0.01 as assessed by unpaired *t* test).

**Figure 12:**
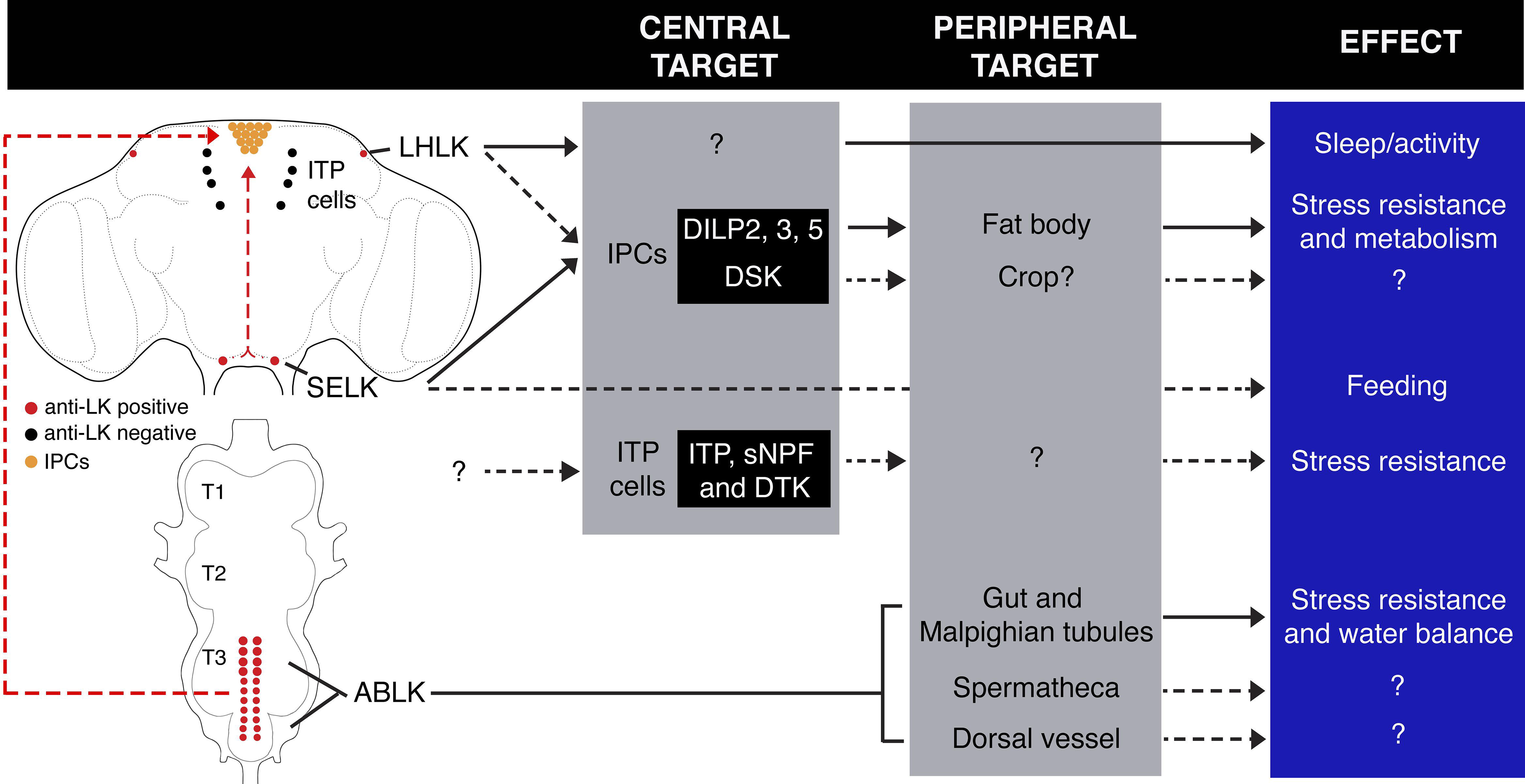
*Lk* signaling scheme. LK signaling scheme showing the location of all LK neurons, identified neurons downstream of LK neurons, target tissues and their effects. Dashed arrows indicate probable links that need to be functionally validated. DSK, drosulfakinin; sNPF, short neuropeptide F; DTK, tachykinin.

## Discussion

In this study we established the role of LK signaling in orchestrating behavioral and physiological homeostasis in *Drosophila*. More specifically, we determined a set of effects caused by loss of LK signaling, which indicates that this neuropeptide regulates physiology related to water homeostasis and metabolism, as well as associated stress, locomotor activity and metabolic rate. We suggest that LK signaling regulates post-feeding physiology, metabolism and behavior, as this seems to link most of the observed phenotypes observed after peptide and receptor knockdown.

In support of the physiological roles of LK signaling, we show distribution of the *Lkr* expression in cells of the renal tubules and intestine, including the water-regulating rectal pads, as well as in the IPCs, which are known to signal with DILPs to affect feeding, metabolism, sleep, activity and stress responses [33–36,38]. *Lkr* is also expressed by another set of brain neurosecretory cells (ipc-1/ipc-2a) known to regulate stress responses by means of three different coexpressed neuropeptides [24].

In the CNS of the adult fly, LK is produced at high levels by a small number of neurons of three major types: two pairs of interneurons in the brain and about 20 neurosecretory cells, ABLKs, in the abdominal ganglia [6,7]. There is mounting evidence that the ABLKs use LK as a hormonal signal that targets peripheral tissues, including the renal tubules [17] and that the brain LK neurons act in neuronal circuits within the CNS [18–20,39]. More specifically, the LHLK brain neurons are part of the output circuitry of the circadian clock in regulation of locomotor activity and sleep suppression induced by starvation [19,20,39] and the SELKs of the subesophageal zone may regulate feeding [18]. In fact we show here that these SELKs have axons that exit through subesophageal nerves known to innervate muscles of the feeding apparatus. We found in this study that the ABLKs display increased calcium activity in response to drinking in desiccated flies, but not during starvation, desiccation or regular feeding. This finding supports a role of ABLKs and hormonal LK in regulation of water balance. These neurons have also been implicated more broadly in control of water and ion homeostasis and in responses to starvation, desiccation and ionic stress [17]. The LHLKs and SELKs did not display changes in calcium signaling under the tested conditions, strengthening the unique function of ABLKs in diuresis.

The regulation of metabolic rate, as determined by measurement of CO_2_ production, is a novel phenotype that we can link to LK signaling. This may be associated with the overall activity of the flies, as suggested by the correlation between activity and CO_2_ levels in our data. Thus, the regulation of activity and metabolic rate might be coordinated by means of the LK neurons.

Using anatomical and experimental strategies, we identified a novel circuit linking LK to insulin signaling. *Lkr* expression was detected in the brain IPCs using two independently generated *GAL4* lines plus single-cell transcriptome analysis. We also observed that *Lk* and *Lkr* mutants displayed increased levels of DILP2 and DILP3 immunoreactivity in the brain IPCs and targeted knockdown of *Lkr* in IPCs increased *DILP3* expression. Associated with this we found that *Lkr*-RNAi targeted to IPCs increased resistance to desiccation. However, using the *trans*-Tango method for anterograde trans-synaptic labeling [30], we could not demonstrate direct synaptic inputs to IPCs from LK neurons. The LHLKs did not yield any detectable signal; however, the *Lk*-GAL4 line displayed very weak expression in the LHLKs. The SELKs drove postsynaptic marker signal in sets of neurons in the SEG, some of which have processes impinging on the IPCs. These findings suggest that SELKs form no conventional synaptic contacts with IPCs, but paracrine LK signaling to these neurons is not excluded since the two sets of neurons have processes in close proximity in the tritocerebrum and the subesophageal zone. Nonsynaptic paracrine signaling with neuropeptides has been well established in mammals (see [40–42]) and is likely to occur also in insects. Alternatively, the LK input to IPCs could occur systemically at the peripheral axon terminations of the IPCs after hormonal release from ABLKs. Whether paracrine or hormonal, LK appears to regulate the IPCs and transcription and release of DILPs. Thus, some phenotypes seen after the global knockdown of LK and its receptor are likely to arise via secondary effects on insulin signaling, suggesting another layer of regulatory control whereby LK-modulation of DILP production and release could affect metabolism, stress responses and longevity [reviewed by [38,43,44]]. Our findings, therefore, add LK as yet another regulator of the *Drosophila* IPCs, which have previously been shown to be under the regulation of several other neuropeptides and neurotransmitters [reviewed in [38,43]]. It is noteworthy that at the levels of both transcription and presumed release the LK effect on IPCs is selective, affecting DILP2, DILP3 and *DILP3* only.

We suggest that LK signaling regulates post-feeding physiology and behavior seen in the mutants as reduced metabolic rate and locomotor activity, diminished PER, and reduced diuresis, as well as increased resistance to starvation and desiccation. Our data also indicate that in wild type flies LK triggers release of IPC-derived DILPs that are required for post-feeding metabolism and satiety, and it acts on other cells to induce diuresis, and to increase activity (especially evening activity) and metabolic rate. An orchestrating role of LK signaling requires that the three types of LK neurons communicate with each other or are under simultaneous control by common sets of regulatory neurons. Alternatively, all the LK neurons could possess endogenous nutrient-sensing capacity whereby they can monitor levels of amino acids or carbohydrates in the organism. There is evidence for nutrient sensing in LHLK neurons [45]. This has also been shown for the DH44, DILP and corazonin expressing brain neurosecretory cells [31,46–48]. Of the LK neurons, only the ABLKs and SELKs exhibit overlapping processes that could support direct communication, so it is more likely that other neurons form the link between this set of neuroendocrine cells. Such neurons are yet to be identified, but it has been shown that all the LK neurons express the insulin receptor, dInR [21]. This may suggest that the LK neurons receive nutrient-related information from insulin-producing cells in the brain or elsewhere.

In conclusion, we found that LK signaling is likely to orchestrate postprandial physiology and behavior in *Drosophila*. Food ingestion is followed by increased insulin signaling, activation of diuresis, increased metabolic rate, and lowered locomotor activity and increased sleep [10,19,31,43]. Flies mutated in the *Lk* and *Lkr* genes display phenotypes consistent with a role in regulation of insulin signaling, metabolic stress responses, diuresis, metabolic rate, and locomotor activity, all part of postprandial physiology.

## Experimental procedures

### Fly lines and husbandry

All fly strains used in this study (Table 1) were reared and maintained at 25°C on enriched medium containing 100 g/L sucrose, 50 g/L yeast, 12 g/L agar, 3ml/L propionic acid and 3 g/L nipagin, unless otherwise indicated. Experimental flies were reared under normal photoperiod (12 hours light: 12 hours dark; 12L:12D). Adult males 6-8 days post-eclosion were used for behavioral experiments. For some imaging experiments females of the same age were also utilized. For *trans*-Tango analysis, flies were reared at 18°C and adult males 2-3 weeks old post-eclosion were used.

For *DILP2>Lkr-RNAi* qPCR, crosses were established in normal food (NutriFly Bloomington formulation) and eggs were laid for 24 hours. After adult eclosion, males were transferred to alternative diets (normal diet described above; high-sugar high-protein: normal diet except with 20% sucrose and 10% yeast; low-sugar high-protein: normal diet except 5% sucrose and 10% yeast). After 5-7 days on these media, heads were dissected for qPCR.

**Table 1:**
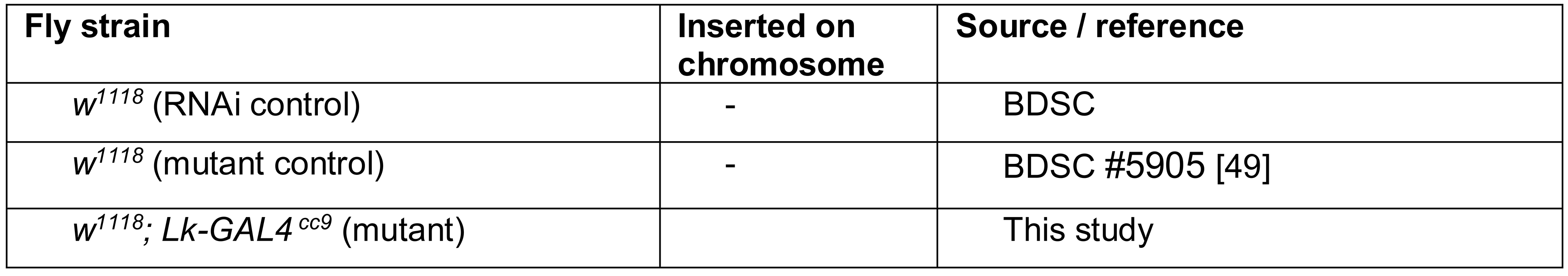
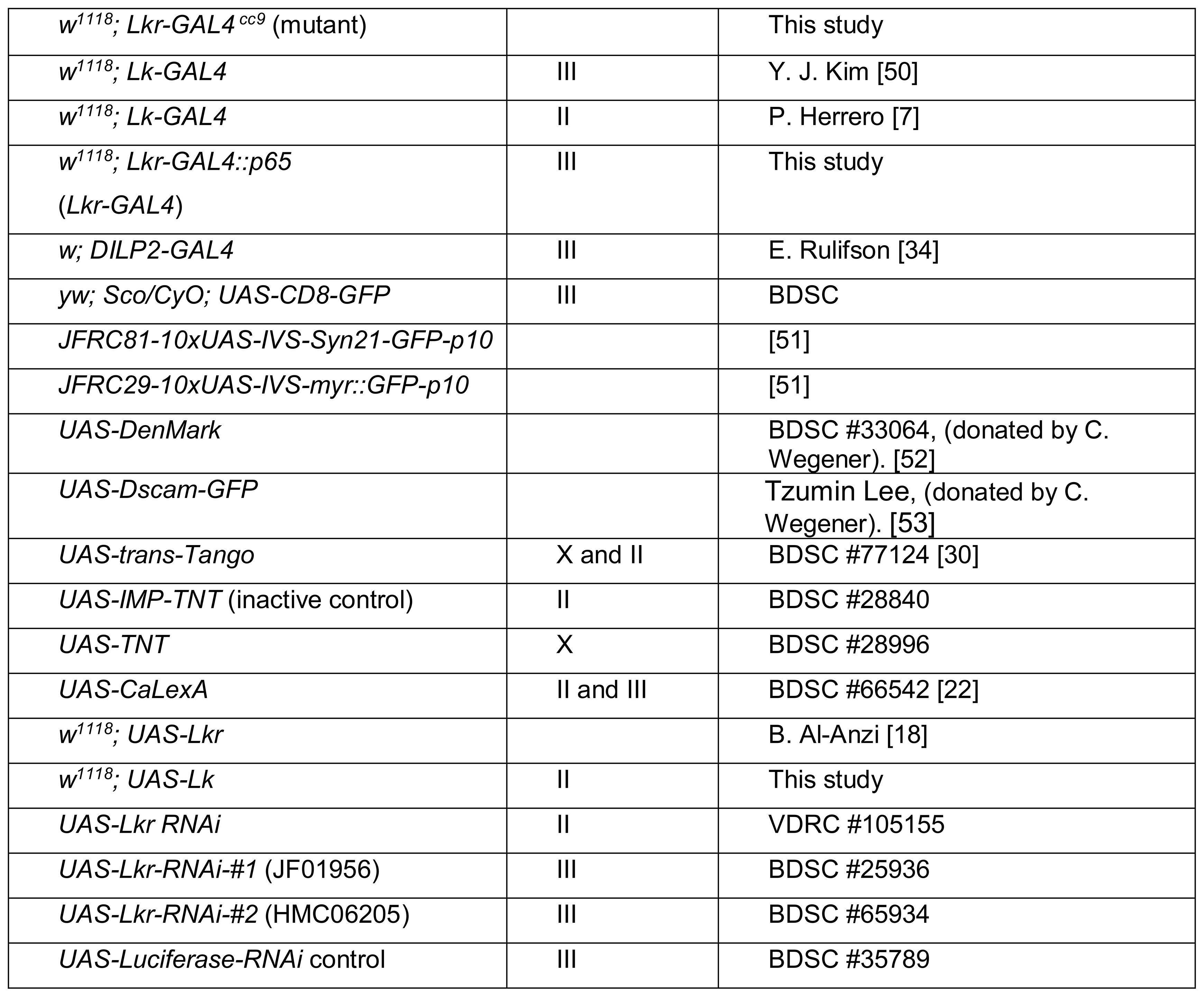
Fly strains used in this study

### Generation of GAL4 knock-in mutants and transgenic lines

*Lk−/−* and *Lkr* −/− were generated using the CRISPR/Cas9 system to induced homology-dependent repair (HDR) using one guide RNA (*Lk*−/−: GATCTTTGCCATCTTCTCCAG and *Lkr*−/−: GTAGTGCAATACATCTTCAG). At gRNA target sites a donor plasmid was inserted containing a GAL4::VP16 and floxed 3xP3-RFP cassette. For *Lk* −/−, the knockin cassette was incorporated immediately following the ATG start site (4bp to 10bp, relative to start site). For *Lkr*−/−, the knock in cassette was incorporated upstream of the ATG start site (−111 bp to −106bp, relative to start site). All lines were generated in the *w*^*1118*^ background. Proper insertion loci for both mutations were validated by genomic PCR. CRISPR gene editing was done by WellGenetics (Taipei City, Taiwan).

To prepare the *Lkr-GAL4::p65* line, recombineering approaches based on previous methods [54] were used (briefly, containing a large genomic BAC with *GAL4::p65* replacing the first coding region of *Lkr*, thereby retaining regulatory flanks and introns). First, a landing-site cassette was prepared: GAL4 and terminator homology arms were amplified from *pBPGUw* [55] and added to the flanks of the marker *RpsL-kana* [56], which confers resistance to kanamycin and sensitivity to streptomycin. *Lkr*-specific arms were added to this landing-site cassette by PCR with the following primers, made up of 50 bases of *Lkr*-specific homology (lower case) plus regions matching the GAL4/terminator sequences:

Lkr-F: tcatatcctcattaggatacacaactaaaactaaaaaacgaaaaagtgtt<u>ATG</u>AAGCTACTGTCTTCTATCGAACAAGC
Lkr-R: tggatgagtcgcgtccccagttgcttgaagggattagagagtatacttacGATCTAAACGAGTTTTTAAGCAAACTCACTCCC

Note the underlined ATG, reflecting the integration of *GAL4* at the *Lkr* initiation site. The PCR product was recombined into bacterial artificial chromosome CH321-16C22 [57] (obtained from Children’s Hospital Oakland Research Institute, Oakland, CA, USA), which contains the *Lkr* locus within 90 kb of genomic flanks. Recombinants were selected on kanamycin. Next, this landing pad was replaced by full-length GAL4::p65+terminators amplified from *pBPGAL4.2::p65Uw* [58], and recombinants were screened for streptomycin resistance. Recombination accuracy was confirmed by sequencing, and the construct was integrated into *attP40* by Rainbow Transgenic Flies (Camarillo, CA, USA).

### RT-qPCR

To quantify *Lk* and *Lkr* transcript levels in mutant flies, the following method was used. Briefly, ten or more fed flies were flash frozen for each sample. Total RNA was extracted from whole flies using RNeasy Tissue Mini kit (QIAGEN) according to the manufacturer’s protocol. RNA samples were reverse transcribed using iScript (Biorad), and the subsequent cDNA was used for real-time RT-qPCR (Biorad CFX96^TM^, SsoAdvanced^TM^ Universal SYBR® Green Supermix qPCR Mastermix Plus for SYBRGreen I) using 1.7ng of cDNA template per well and a primer concentration of approximately 300nM. The primers used are listed in Table 2. Triplicate measurements were conducted for each sample.

To quantify *DILP2, 3* and *5* transcript levels following *DILP2*>*Lkr RNAi*, the following method was used. *DILP2-GAL4* and *UAS-RNAi* animals (*Lkr-RNAi-#1* and −#*2*, plus *UAS-Luciferase-RNAi* as a control for effects of genetic background and RNAi induction) were mated and allowed to lay eggs for 24 hours in vials containing normal food; adult males from these crosses were then transferred to vials of normal food or high-sugar, high-protein or low-sugar high-protein diet. After 7 days, heads were dissected on ice into extraction buffer, and RNA was extracted with the Qiagen RNeasy Mini kit (#74106) with RNase-free DNase treatment (Qiagen #79254). cDNA was prepared using the High-Capacity cDNA Reverse Transcription Kit with RNase Inhibitor (ThermoFisher #4268814), and qPCR was performed using the QuantiTect SYBR Green PCR Kit (Fisher Scientific #204145) and an Mx3005P qPCR system (Agilent Technologies). Expression levels were normalized against RpL32 (Rp49), whose levels have been determined to be stable under dietary modification [32,59]. The primers used are listed in Table 2. Samples were prepared in four biological replicates of 10 heads each, and each biological replicate was assayed in two technical replicates.

**Table 2:**
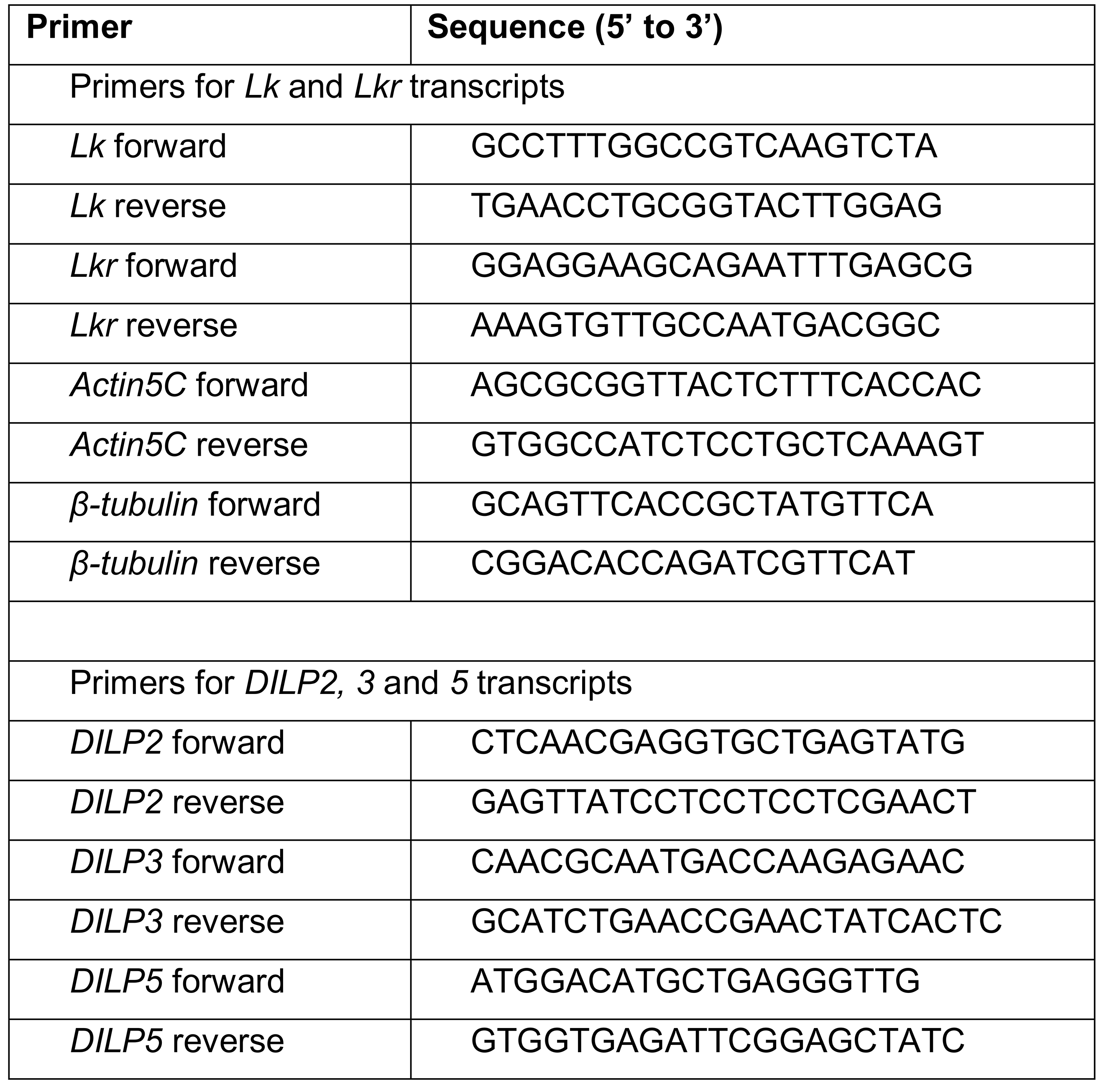
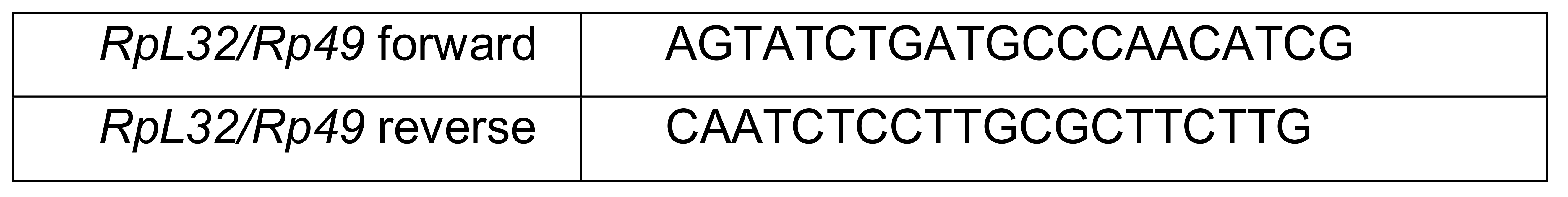
Primers used for qPCR

### Immunohistochemistry and imaging

Immunohistochemistry for *Drosophila* larval and adult tissues was performed as described earlier [17,60]. Briefly, tissues were dissected in phosphate buffered saline (PBS) and fixed in 5% ice-cold paraformaldehyde (2 hours for larval samples and 3.5 – 4 hours for adults). Samples were then washed in PBS and incubated for 48 hours at 4°C in primary antibodies diluted in PBS with 0.5% Triton X (PBST) (Table 3). Samples were thereafter washed with PBST and incubated for 48 hours at 4°C in secondary antibodies diluted in PBST (Table 3). Following this incubation, some samples (peripheral tissues) were incubated with rhodamine-phalloidin (1:1000; Invitrogen) and/or DAPI as a nuclear stain (1:1000; Sigma) diluted in PBST for 1 hour at room temperature. Finally, all samples were washed with PBST and then PBS, and mounted in 80% glycerol. An alternate procedure was used for the adult gut to prevent tissues from rupturing. Briefly, intestinal tissues (proventriculus, crop, midgut, hindgut and MTs) were fixed at room temperature for 2 hours, washed in PBS, incubated in rhodamine-phalloidin for 1 hour and washed in PBST and then PBS before mounting. Samples were imaged with a Zeiss LSM 780 confocal microscope (Jena, Germany) using 10X, 20X or 40X oil immersion objectives. Images for the whole fly, proboscis and wing were captured using a Zeiss Axioplan 2 microscope after quickly freezing the fly at −80°C. Cell fluorescence was measured as described previously [17]. Confocal and fluorescence microscope images were processed with Fiji [61] for projection of z-stacks, contrast and brightness, and calculation of immunofluorescence levels.

**Table 3:**
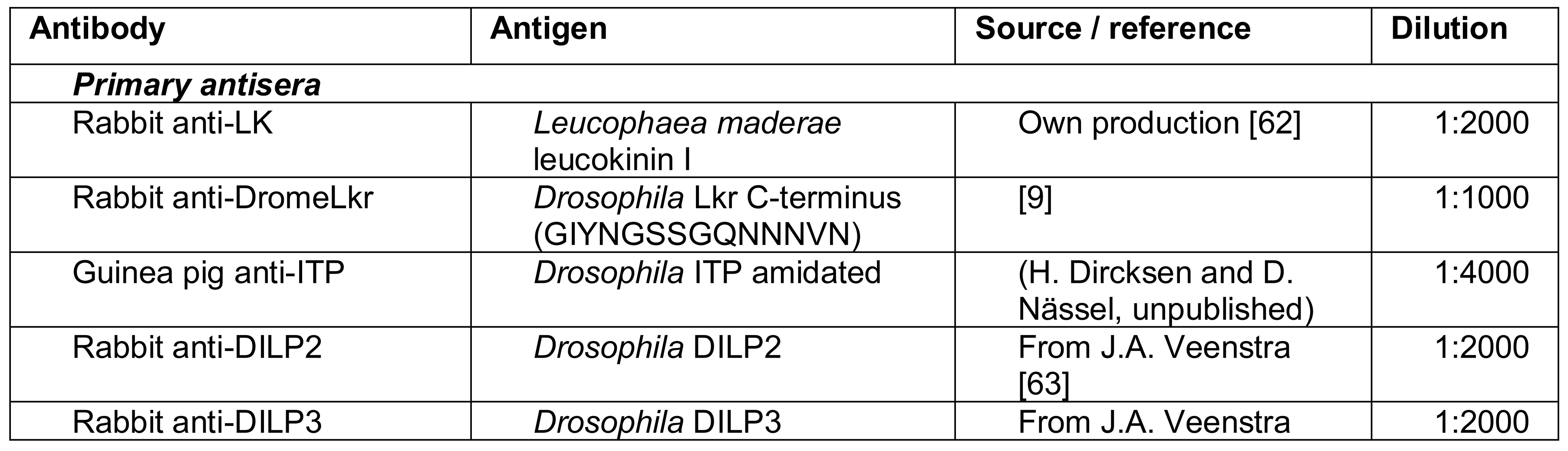
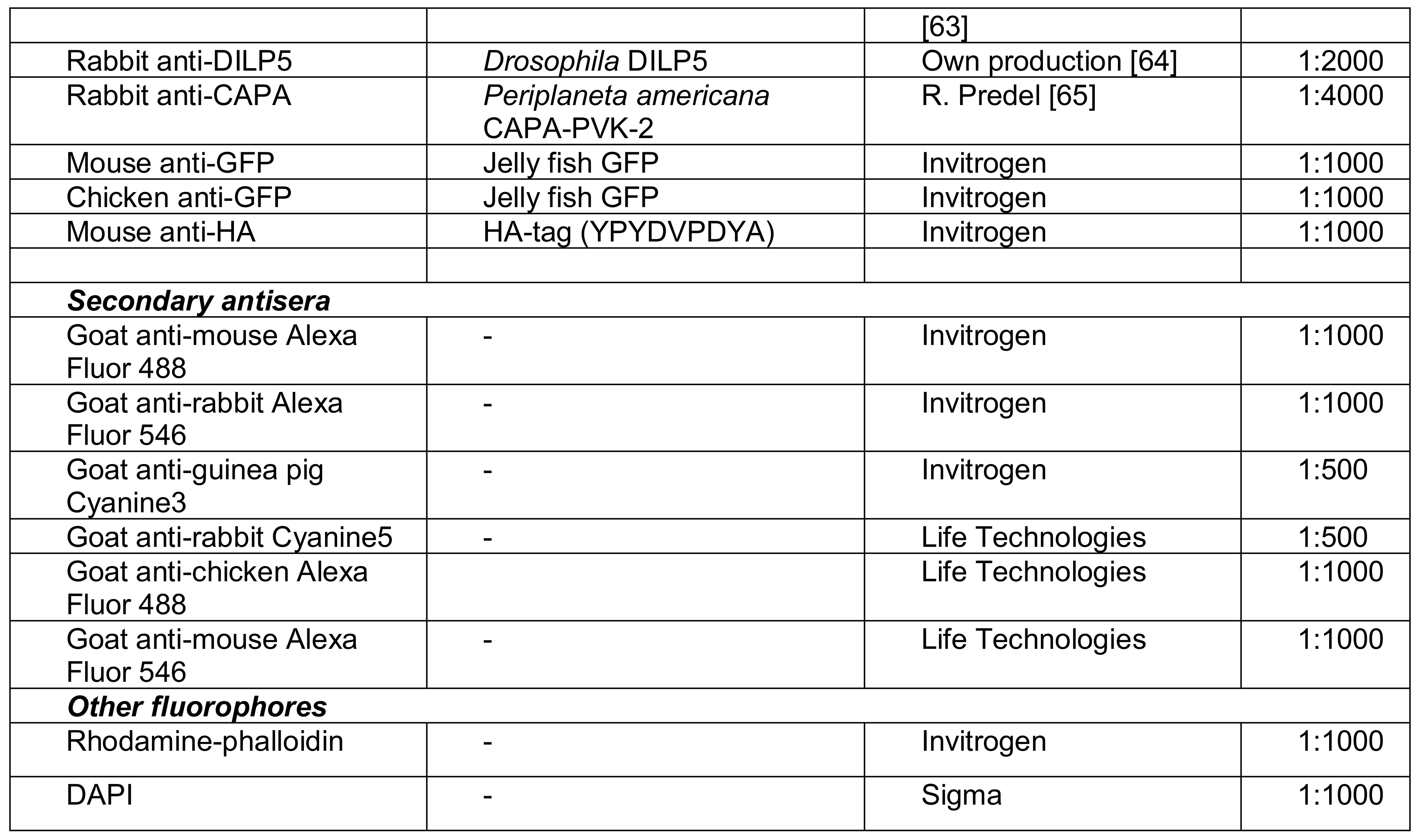
Antibodies used for immunohistochemistry

### Calcium activity in LK neurons

Calcium activity of LK neurons following various stresses was measured using the CaLexA (<u>Cal</u>cium-dependent nuclear import of <u>LexA</u>) technique [22]. Briefly, 6-8-day-old males were either transferred to a vial containing nothing (desiccation), a vial containing aqueous 1% agar (starvation) or a vial containing artificial diet (normal food) and incubated for 16 hours. In addition, one set of flies were desiccated for 13 hours and then transferred to a vial containing 1% agar (re-watered). Following this period, the flies were fixed, dissected brains processed for immunohistochemistry and the GFP fluorescence was quantified as described above.

### Stress-resistance assays

To assay for survival under desiccation (dry starvation) and starvation, flies were kept in empty vials and vials containing 5 ml of 0.5% aqueous agarose (A2929, Sigma-Aldrich), respectively. Four biological replicates and 3 technical replicates for each biological replicate were performed for each experiment. For each technical replicate, 15 flies were kept in a vial and their survival was recorded every 3 to 6 hours until all the flies were dead. The vials were placed in incubators at 25°C under normal photoperiod conditions (12L:12D).

### Water-content measurements

For water content measurements, 15 flies per replicate (4 biological replicates) were either frozen immediately on dry ice or desiccated as above for 9 hours and then frozen. The samples were stored at −80°C until use. To determine their wet weight, flies were brought to room temperature and their weight was recorded using a Mettler Toledo MT5 microbalance (Columbus, USA). The flies were then dried for 24-48 hours at 60°C before recording their dry weight. The water content of the flies was determined by subtracting dry weight from wet weight.

### Blue dye feeding assay

Short-term food intake was measured as previously described [66]. Briefly, flies were starved for 24 hours on 1% agar (Fisher Scientific) or maintained on standard fly food. At ZT0, flies were transferred to food vials containing 1% agar, 5% sucrose, and 2.5% blue dye (FD&C Blue Dye No. 1, Spectrum). Following 30 minutes of feeding, flies were flash frozen on dry ice and four flies per sample were homogenized in 400 μL PBS (pH 7.4, Fisher Scientific). Color spectrophotometry was used to measure absorbance at 655 nm in a 96-well plate reader (Millipore, iMark, Bio-Rad). Baseline absorbance was determined by subtracting the absorbance measured in non-dye fed flies from each experimental sample.

### Proboscis extension reflex

Flies were collected and placed on fresh food for 24 hours, then starved for 24 hours in vials containing 1% agar. Flies were then anaesthetized under CO_2_, and their thorax and wings were glued with nail polish to a microscopy slide, leaving heads and legs unconstrained. Following 1-hour recovery in a humidified chamber, the slide was mounted vertically under the dissecting microscope (SM-3TX-54S, AmScope) and proboscis extension reflex (PER) was observed. PER induction was performed as described previously [67]. Briefly, flies were satiated with water before and during experiments. Flies that did not water satiate within 5 minutes were excluded from the experiment. A 1 ml syringe (Tuberculin, BD&C) with an attached pipette tip was used for tastant (sucrose) presentation. Tastant was manually applied to tarsi for 2-3 seconds 3 times with 10 second inter-trial intervals, and the number of full proboscis extensions was recorded. Tarsi were then washed with distilled water between applications of different concentrations of sucrose (0.1, 1.0, 10, and 100 mM) and flies were allowed to drink water during the experiment *ad libitum*. Each fly was assayed for response to tastants. PER response was calculated as a percentage of proboscis extensions to total number of tastant stimulations to tarsi.

### Activity and metabolic rate

Activity and metabolic rate (MR) was simultaneously recorded using the setup described earlier [23]. Briefly, MR was measured at 25°C through indirect calorimetry, measuring CO_2_ production of individual flies with a CO_2_ analyzer (LI-7000, LI-COR). Baseline CO_2_ levels were measured from an empty chamber, alongside five behavioral chambers, each measuring the CO_2_ production of a single male fly. The weight of a group of 10 flies was used to normalize metabolic rate since *Lk* mutants weighed significantly more than control *w*^*1118*^ flies. Flies were anesthetized using CO_2_ for sorting and allowed 24 hours acclimation before the start of an experiment. Flies were placed in glass tubes that fit a custom-built *Drosophila* Locomotor Activity Monitor (Trikinetics, Waltham, MA), containing a single food tube containing 1% agar plus 5% sucrose with green food coloring (McCormick). Locomotor activity data was calculated by extracting 10 minute activity periods for 24 hours using a custom generated Python program. CO_2_ output was measured by flushing air from each chamber for 75 seconds providing readout of CO_2_ accumulation over the 10-minute period. This allowed for the coordinate and simultaneous recordings of locomotor activity and metabolic rate.

### Locomotor Activity

*Drosophila* activity monitoring system (DAMS; Trikinetics, Waltham, MA) detects activity by monitoring infrared beam crossings for each animal. These data were used to calculate locomotor activity using the *Drosophila* Sleep Counting Macro [68]. Flies were anaesthetized under CO_2_ and loaded into DAMS tubes containing standard fly food for acclimation. After 24 hours acclimation in DAMS tubes with food, baseline activity was measured for 24 hours. Tubes were maintained in a 25°C incubator with 12:12 LD cycles.

### Mining public datasets for expression of genes

*Lkr* distribution in various tissues was determined by mining the FlyAtlas database [27]. *Lkr* expression in the different regions of the gut and its cell types was obtained using Flygut-*seq* [28]. A single-cell transcriptome atlas of the *Drosophila* brain was mined using SCope (http://scope.aertslab.org) to identify genes coexpressed with *Lkr* [29].

### Statistical analyses

The experimental data are presented as means ± s.e.m. Unless stated otherwise, one-way analysis of variance (ANOVA) followed by Tukey’s multiple comparisons test was used for comparisons between three genotypes and an unpaired *t* test was used for comparisons between two genotypes. All statistical analyses were performed using GraphPad Prism with a 95% confidence limit (*p* < 0.05). Survival and stress curves were compared using Mantel–Cox log-rank test.

## Acknowledgements

We are grateful to the Bloomington *Drosophila* Stock Center, the Vienna *Drosophila* Resource Center and Drs. Julian Dow, Pilar Herrero, Reinhard Predel, Patricia Pietrantonio and Jan A. Veenstra, for providing flies and reagents. Stina Höglund and the Imaging Facility at Stockholm University (IFSU) are acknowledged for maintenance of the confocal microscopes. We thank Dr. Yiting Liu for providing images of IPC dendrites and Dr. Wouter van der Bijl for assistance in creating Figure 11. This work was supported by a grant from the European Commission Horizon 2020 (Research and Innovation Grant 634361) to D.R.N. and National Institute of Health award R01NS085152 to A.C.K, and Danish Council for Independent Research, Natural Sciences grant 4181-00270 to Kim F. Rewitz. The *Lkr-GAL4::p65* line was generated by M.J.T. at HHMI Janelia Research Campus in the lab of J. Truman.

## Captions for supplementary figures

**Supplementary Table 1:** p-values for the proboscis extension reflex data in Figure 5. p-values below 0.05 have been highlighted in grey. Wilcoxon Rank-Sum was used for comparison between two genotypes, while Kruskal-Wallis with Steel-Dwass post-hoc test was used for two or more genotypes. These tests were performed at each concentration independently.

**Figure S1: Total activity (measured using DAMS) of *Lk* and *Lkr* mutants.** Total locomotor activity of single flies measured over 24 hours is lowered for homozygous and heterozygous **(A)** *Lk* and **(B)** *Lkr* mutants. The activity was monitored using a standard *Drosophila* Activity Monitor (DAMS). (*** p < 0.001, **** p < 0.0001, as assessed by oneway ANOVA).

**Figure S2: The *Lk-GAL4*^*CC*9^ drives GFP expression in the adult CNS.** *Lk-GAL4*^*CC*9^ drives GFP (*pJFRC81-10xUAS-Syn21-myr::GFP-p10*) expression in the adult **(A)** brain and **(B)** ventral nerve cord (VNC). SELK, subesophageal LK neurons; ABLK, abdominal LK neurons. *Lk-GAL4*^*CC*9^ also drives GFP expression in four pairs of neurons in the brain (indicated by the white box). **(C)** These four pairs of neurons display very weak LK-immunoreactivity and are positive for ion transport peptide-immunoreactivity. GFP expression also colocalizes with anti-LK staining in the SELKs and lateral horn LK neurons (LHLK). **(D)** *Lk-GAL4*^*CC*9^ drives GFP expression in ABLKs (labeled with anti-LK antiserum) in the VNC.

**Figure S3: *Lk-GAL4*^*CC*9^ and *Lkr-GAL4*^*CC*9^ drive GFP expression in the larval CNS. (A)** *Lk-GAL4*^*CC*9^ drives GFP (*pJFRC81-10xUAS-Syn21-myr::GFP-p10*) expression in neurosecretory cells in the larval brain and ventral nerve cord (NVC). **(B)** *Lkr-GAL4*^*CC*9^ drives GFP (*UAS-mCD8GFP*) expression in larval CNS. Note the GFP expression in motor neurons in the VNC.

**Figure S4: The *Lkr-GAL4*^*CC*9^ drives GFP expression in adult peripheral tissues.** *Lkr-GAL4*^*CC*9^ drives GFP (*pJFRC81-10xUAS-Syn21-myr::GFP-p10*) expression in the adult **(A)** dorsal vessel and peripheral neurons (indicated by an arrow), **(B)** legs, **(C)** proboscis and **(D)** wings. Note the expression of *Lkr* in nerve fibers closely associated with the anti-LK immunostaining in **(A)**.

**Figure S5: The *Lkr-GAL4*^*CC*9^ drives GFP expression in larval gut and Malpighian tubules.** *Lkr-GAL4*^*CC*9^ drives GFP (*pJFRC81-10xUAS-Syn21-myr::GFP-p10*) expression in the larval **(A)** gut, **(B)** gastric caeca and anterior midgut, **(C)** midgut and **(D)** anti-DromeLkr expressing stellate cells in Malpighian tubules. Nuclei in all the preparations have been stained with DAPI (blue).

**Figure S6: The *Lkr-GAL4* drives GFP expression in gut and Malpighian tubules.** *Lkr-GAL4* drives GFP (*pJFRC29-10xUAS-myr::GFP-p10*) expression in **(A)** the larval stellate cells of Malpighian tubules, **(B)** larval hindgut and **(C-E)** adult stellate cells (labeled with anti-DromeLkr antiserum). Note that the adult stellate cells can be (**C**) cuboidal or (**D**) star-shaped (indicated by an arrow).

**Figure S7: *Lkr-GAL4* drives GFP (*UAS-mCD8-GFP*) expression in larval and adult CNS. (A)** *Lkr-GAL4* drives GFP expression in several neurons of the larval CNS, including a pair of abdominal Lk neurons stained with anti-Lk antiserum (indicated with a white arrow). In adults, *Lkr-GAL4* drives GFP expression in **(B)** T1 and T2 thoracic neuromeres and, **(C)** T3 thoracic neuromere.

**Figure S8: The *Lkr-GAL4*^*CC*9^ drives GFP expression in the adult CNS.** *Lkr-GAL4*^*CC*9^ drives GFP (*UAS-mCD8GFP*) expression in **(A)** the brain and **(B)** ventral nerve cord. The inset in **(A)** represents a smaller Z-stack which shows GFP expression in the fan-shaped body. These preparations were counterstained with anti-nc82 antiserum. **(C)** *Lkr-GAL4*^*CC*9^ drives GFP (*pJFRC81-10xUAS-Syn21-myr::GFP-p10*) expression in neurons of the abdominal ganglia that do not express LK.

**Figure S9: Anatomical interactions between LK and CAPA/hugin signaling. (A)** Expression of *trans*-Tango components [30] using *Lk*-GAL4 generates a post-synaptic signal (labeled with anti-HA antibody) in the tritocerebrum and pars intercerebralis which does not colocalize with CAPA/hugin axons (labeled with anti-CAPA antibody). **(B)** Higher magnification of the subesophageal ganglion showing the pre-synaptic and post-synaptic signals and the lack of colocalization with anti-CAPA staining.

**Figure S10: The processes of IPCs in pars intercerebralis and tritocerebrum/ subesophageal zone have dendrite properties.** Using dendrite-directed UAS constructs, fluorescent labeling can be seen in IPC processes in pars intercerebralis and tritocerebrum/subesophageal zone, shown in inverted images. **(A)** *DILP2-GAL4* driven *Dscam-GFP* and **(B)** *DILP2-GAL4* driven *DenMark-RFP*. These images were kindly provided by Dr. Yiting Liu.

**Figure S11: DILP5 levels are unaltered in *Lk* and *Lkr* mutants. (A)** *Lk* and *Lkr* homozygous mutants do not display any difference in DILP5 immunoreactivity in insulin-producing cells (IPCs) of the adult brain. **(B)** Fluorescence intensity measurement of IPCs shows no difference in DILP5 immunoreactivity in *Lk* and *Lkr* mutant flies compared to control flies. CTCF, corrected total cell fluorescence.

**Figure S12: *Lkr* knockdown in insulin-producing cells.** Knockdown of *Lkr* in IPCs has **(A)** no effect on starvation, results in **(B)** increased survival under desiccation and **(C)** an increase in dry weight. (* p < 0.05 as assessed by Log-rank (Mantel-Cox) test for **(B)**, and * p < 0.05 and ** p < 0.01 for **(C)** as assessed by one-way ANOVA followed by Tukey’s multiple comparisons test).

